# A novel major output target for pheromone-sensitive projection neurons in male moths

**DOI:** 10.1101/804922

**Authors:** Xi Chu, Stanley Heinze, Elena Ian, Bente G. Berg

## Abstract

The male-specific macroglomerular complex (MGC) in the moth antennal lobe contains circuitry dedicated to pheromone processing. Output neurons from this region project along three parallel pathways, the medial, mediolateral, and lateral tracts. The MGC-neurons of the lateral tract are least described and their functional significance is unknown. We used mass-staining, calcium imaging, and intracellular recording/staining to characterize the morphological and physiological properties of these neurons in *Helicoverpa armigera*. All lateral-tract MGC neurons targeted the *column*, a small region within the superior intermediate neuropil. We identified this region as the major converging site for lateral-tract neurons responsive to pheromones and plant-odors. The lateral-tract MGC-neurons consistently responded with a faster onset than the well-described medial-tract neurons. Different from the medial-tract MGC neurons encoding odor quality important for signal identification, those in the lateral tract seem to convey a robust and rapid, but fixed signal – potentially important for fast control of hard-wired behavior.

## Introduction

Pheromones are chemical signals serving in social and sexual communication between individuals of distinct species throughout the animal kingdom. While the peripheral mechanisms for pheromone detection are well described in many species, knowledge about central processing principles of these key sensory stimuli remains incomplete. Noctuid moths contain numerous prime examples of species with highly specific pheromone communication combined with exquisite sensitivity. Among the most intensively investigated are several species of the subfamily Heliothinae (Lepidoptera: Noctuidae) (Cho et al., 2008, Fitt, 1989), including some of the world’s most detrimental crop pests. In contrast to the also intensively studied, domesticated silk moth, *Bombyx mori*, heliothine moths are *flying* species utilizing pheromones to communicate over long distances. The males recognize minute amounts of the female-produced pheromones via highly sensitive sensory neurons housed in specialized sensilla on the antennae.

In most moths, the male-specific sensory neurons project directly to a distinct region in the antennal lobe, the macroglomerular complex (MGC). This pathway is dedicated to process input about female-produced compounds and is present in addition to the general olfactory circuit involving the usually more numerous ordinary AL glomeruli. The species used in this study, *Helicoverpa armigera*, utilizes two pheromone components: *cis-*11-hexadecenal (Z11-16:Al) as primary component and *cis-*9-hexadecenal (Z9-16:Al) as secondary component (Kehat and Dunkelblum, 1990). The MGC of this species contains three glomeruli, named cumulus, dorsomedial anterior (dma), and dorsomedial posterior (dmp; Skiri et al., 2005, Zhao et al., 2016a). Originating from the sensory neurons in specific antennal sensillae, signals resulting from the two pheromone components as well as one behavioral antagonist, *cis*-9-tetradecenal (Z9-14:Al), are received by the cumulus, dmp, and dma, respectively (see Figure 7 in Wu et al., 2015).

In the antennal lobe, all sensory axons make synaptic contacts with local interneurons and projection neurons. The latter cells carry odor information to higher integration centers in the protocerebrum via several parallel antennal-lobe tracts (ALTs). The three main ALTs, the medial ALT, the mediolateral ALT, and the lateral ALT, connect the antennal lobe with the calyces of the mushroom bodies (MB) and the lateral protocerebrum (mostly lateral horn), which constitute the two most prominent higher-order olfactory projection areas across insects (Homberg et al., 1988, Seki et al., 2005, Rø et al., 2007, Ito et al., 2014). A third area is targeted by a significant proportion of lateral-tract projection neurons and is embedded in the superior intermediate protocerebrum (SIP, see Ito et al., 2014), occupying the space in between the anterior optic tubercle (AOTU) and the MB vertical lobe. Although it was discovered in the hawk moth *Manduca sexta* (Homberg et al., 1988), its prominence as a major projection region was pointed out in the heliothine moth, where it was termed the *column* (Ian et al., 2016).

Similar to insects, an arrangement of parallel olfactory tracts and projection areas is also found in vertebrates, such as fish (Hamdani and Døving, 2007) and mammals (Mori, 2016, Kauer, 1991). In fish, each of three tracts carries a different category of olfactory information, i.e. social cues, pheromones, and food odors (Hamdani and Døving, 2007). Contrary, in insects, particularly in moths, each of the three main ALTs is formed by axons of projection neurons originating from both the MGC and the ordinary glomeruli (Homberg et al., 1988, Zhao et al., 2014, Kanzaki et al., 2003). Thus, different categories of olfactory cues are processed in all ALTs. Rather than encoding different stimulus categories, the parallel tracts in insects are likely to transmit information about different features (e.g. concentrations) of the same odor (reviewed by Galizia and Rossler, 2010). Due to the limited number of involved odors and their high relevance to behavior, examining pheromone processing provides a low-dimensional model to illuminate the functional differences across parallel pathways.

To reveal these differences, functional and anatomical work on neurons in all ALTs is required. However, studies on individual MGC projection neurons in moths have focused almost exclusively on uni-glomerular medial-tract neurons. These previous reports include studies of heliothine moths, e.g. *Heliothis virescens*, *Helicoverpa zea*, and *Helicoverpa assulta* (Christensen et al., 1991, Christensen et al., 1995, Vickers et al., 1998, Zhao and Berg, 2010, Zhao et al., 2014), as well as *Manduca sexta* (Christensen and Hildebrand, 1987, Kanzaki et al., 1989, Hansson et al., 1991), *Agrotis ipsilon* (Jarriault et al., 2009), and *Bombyx mori* (Kanzaki et al., 2003, Seki et al., 2005). Only a recent publication on two heliothine species reported individual pheromone neurons passing along ALTs other than the medial (Lee et al., 2019). Thus, our understanding of pheromone processing across parallel pathways in moths is still very limited. We therefore investigated morphological and physiological properties of male-specific projection neurons confined to the second prominent ALT, the lateral tract. By using intracellular recordings combined with staining, calcium imaging, and mass labeling of neurons, we found that all identified lateral-tract MGC neurons projected to the column. This convergence of lateral-tract axons within a single, restricted region was distinct from the more widespread medial-tract MGC terminals in the superior lateral protocerebrum, which have been suggested to form patterns according to behavioral significance (Zhao et al., 2014). In addition, temporal dynamics and tuning characteristics of the lateral-tract MGC neurons were distinct from those confined to the medial tract. Finally, a substantial proportion of lateral-tract projection neurons originating from the ordinary glomeruli also terminated in the column, exposing this region as a site for convergence for sensory information from the pheromone and plant odor sub-systems. Taken together, the results presented here suggest that the morphologically distinct types of MGC neurons passing in the lateral ALT play other roles than the medial-tract neurons. Particularly, the lateral-tract neurons convey a more direct and pre-coded signal, which might be advantageous for fast initiation of stereotypical plume-tracking flight behavior.

## Results

### Outline of the male Helicoverpa armigera brain

To provide a general anatomical framework for interpreting the functional and morphological information from odor processing neurons we have generated 3D reconstructions of three representative male *H. armigera* brains. Based on synapsin-labeled whole-mount preparations, we segmented all identifiable major neuropils (Figure 1). This 3D brain atlas includes 25 separate neuropils (23 paired and 2 unpaired) and is thus the most detailed representations of any moth brain to date (interactive viewing and download at: www.insectbrain.org; https://hdl.handle.net/20.500.12158/SIN-0000020.2). While particular attention was paid to the reconstruction of olfactory brain regions, such as the antennal lobes and the components of the mushroom body, we also reconstructed all optic lobe neuropils (lamina, medulla, lobula, lobula plate and accessory medulla), as well as the sub-compartments of the anterior optic tubercle (upper unit, lower unit and nodular unit), the lateral complex (lateral accessory lobe, bulb and gall), and the central complex (fan-shaped body, ellipsoid body, protocerebral bridge, noduli, and posterior optic tubercle). Given the unclear boundaries within the remaining major parts of the brain (superior, inferior, ventrolateral, ventromedial, supraesophageal and subesophageal neuropils, as well as the lateral horn), we did not segment these regions separately, but rather used the below described dye fills to highlight relevant compartments in these ‘unstructured’ parts of the brain.

**Figure 1.**
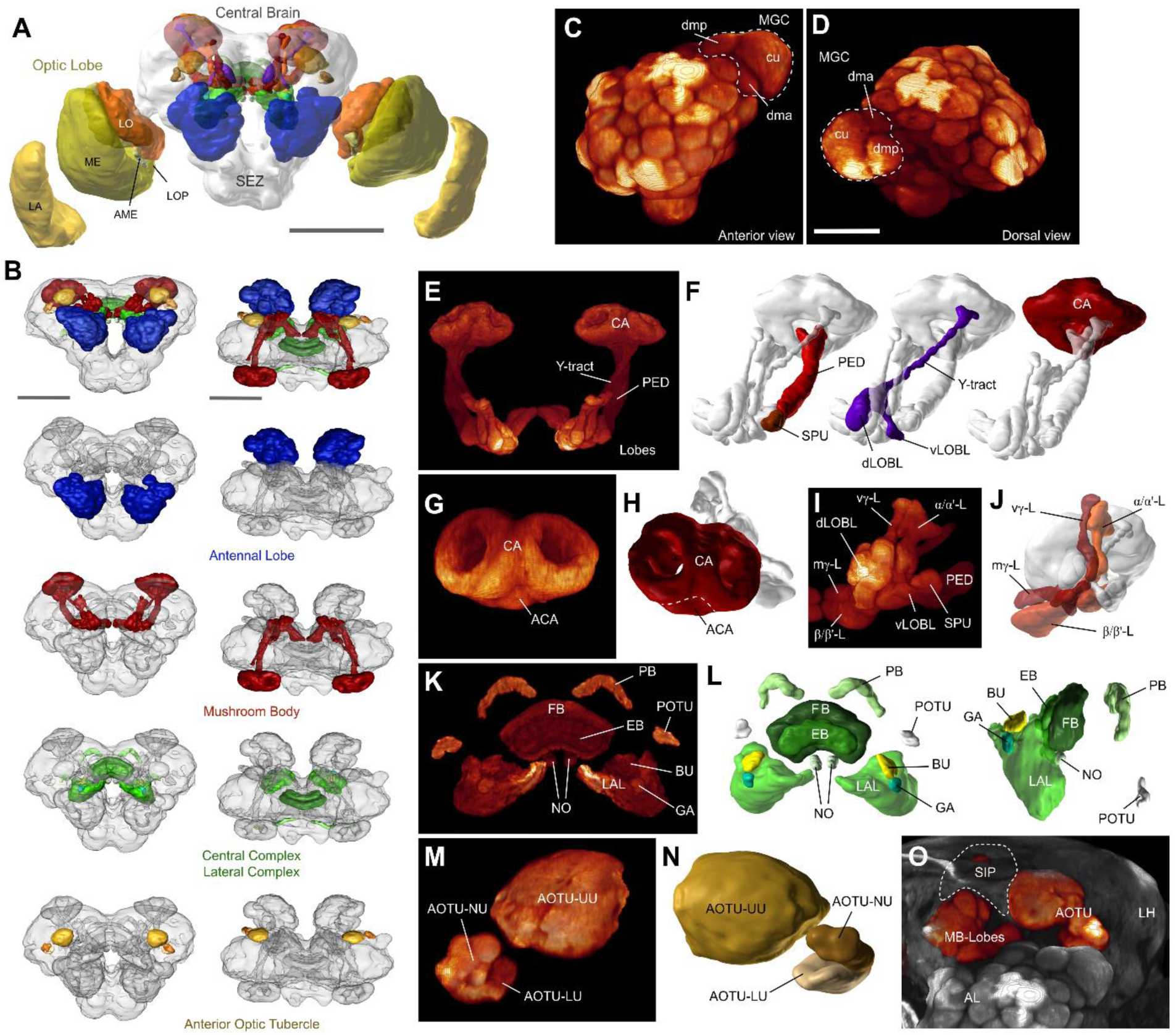
The layout of the *Helicoverpa armigera* brain. **(A)** Three-dimensional reconstruction of an anti-synapsin labeled male brain. Shown are polygonal surface models based on segmented confocal image stacks. LA, lamina; ME, medulla; LO, lobula; LOP lobula plate; AME, accessory medulla; SEZ, subesophageal zone. Scalebar: 500 µm. (**B)** The main defined neuropils of the central brain. *From top to bottom:* All defined central brain neuropils (colored) shown together with the continuous mass of undefined regions (grey); antennal lobes; mushroom bodies; central and lateral complex; anterior optic tubercle. Scalebar: 300 µm (**C-D**) Volume rendering of one male antennal lobe (anti-synapsin labeling), highlighting the location and composition of the macroglomerular complex (MGC); anterior view (**C**), and dorsal view (**D**). Scalebar: 100 µm. (**E**) Volume rendering of the mushroom bodies. All remaining neuropils were masked. (**F**) Principal components of the mushroom body besides the main lobe system: Calyx (CA), pedunculus (PED) with spur (SPU), and Y-tract with dorsal and ventral lobelets (dLOBL, vLOBL). (**G**) Volume rendering of the calyx, highlighting the accessory calyx (ACA), and the two fused rings of the main calyx. (**H**) Surface reconstruction of the calyx. (**I-J**) Volume rendering (**I**) and reconstruction (**J**) of the main lobe systems of the mushroom body. vertical gamma lobe (vγ-L), medial gamma lobe (mγ-L), alpha/alpha’-lobes (α/α’-L), and beta/beta’-lobes (β/β’-L). (**K-L**) Volume rendering (**K**) and reconstruction (**L**) of the central complex and lateral complexes. *Left panel* in (**L**): anterio-dorsal view; *right panel:* lateral view. protocerebral bridge (PB), fan-shaped body (FB), ellipsoid body (EB), noduli (NO), posterior optic tubercle (POTU), bulbs (BU), gall (GA), lateral accessory lobe (LAL). (**M-N**) Volume rendering (**M**) and reconstruction (**N**) of the anterior optic tubercle (AOTU) and its three compartments; upper unit (UU), lower unit (LU), and nodular unit (NU). (**O**) Volume rendering of the frontal portion of the central brain, highlighting the relative location of the AOTU, the mushroom body lobes, the antennal lobes (AL), and the approximate locations of the superior intermediate protocerebrum (SIP, dashed line) and the lateral horn (LH). AOTU and mushroom body lobes were rendered separately, using a different colormap.

Overall, the *H. armigera* brain is a typical lepidopteran brain and resembles the general outline of previously described butterflies (Heinze et al., 2012; Montgomery and Ott, 2015, Montgomery et al. 2016) and hawkmoths (el Jundi et al., 2009; Stöckl et al. 2017). Whereas these species rely heavily on vision and therefore possess large regions dedicated to early visual processing, our nocturnal moth has much smaller optic lobes, in line with its olfactory ecology. Additionally, the first processing station for visual information in the central brain, the anterior optic tubercle, is also smaller compared to the day active butterflies.

With respect to olfactory brain areas, a pronounced sexual dimorphism is represented by a three-unit MGC in *H. armigera* males (Zhao et al., 2016b, Skiri et al., 2005). Accordingly, we also identified these three male-specific glomeruli in the individuals examined in the current study. The main second-order olfactory brain region, the mushroom body, was much smaller in our noctuid moths compared to butterflies and hawkmoths, but nevertheless consists of equally elaborate subdivisions.

Finally, the central and lateral complexes appear largely identical to all other lepidopteran insects studied and in fact comprise the same subdivisions as all insects examined to date. With respect to shape and relative size, these regions most closely reflected the layout of the corresponding areas in the Bogong moth (*Agrotis infusa*) and the Turnip moth (*Agrotis segetum*) (de Vries et al., 2017), underlining the high degree of conservation of these central brain centers.

### The column is a major site of convergence for pheromone and general odor processing

To obtain an anatomical overview of the pheromone and the plant-odor pathways in the moth brain, we applied two different fluorescent dyes into the male antennal lobe (n = 5). Micro-ruby was injected into the MGC and Alexa 488 into the ordinary glomeruli (Fig. 2A, B). These injections confirmed previous findings reporting that projection fields of medial-tract neurons that originate in the MGC are not overlapping with those originating from the ordinary glomeruli (Homberg et al., 1988, Zhao et al., 2014, Seki et al., 2005). While medial-tract neurons from ordinary glomeruli targeted large areas of the mushroom body calyx before terminating in the lateral horn, the MGC projections sent collaterals to a restricted area in the inner layer of the calycal cups and ended in the superior lateral protocerebrum.

**Figure 2.**
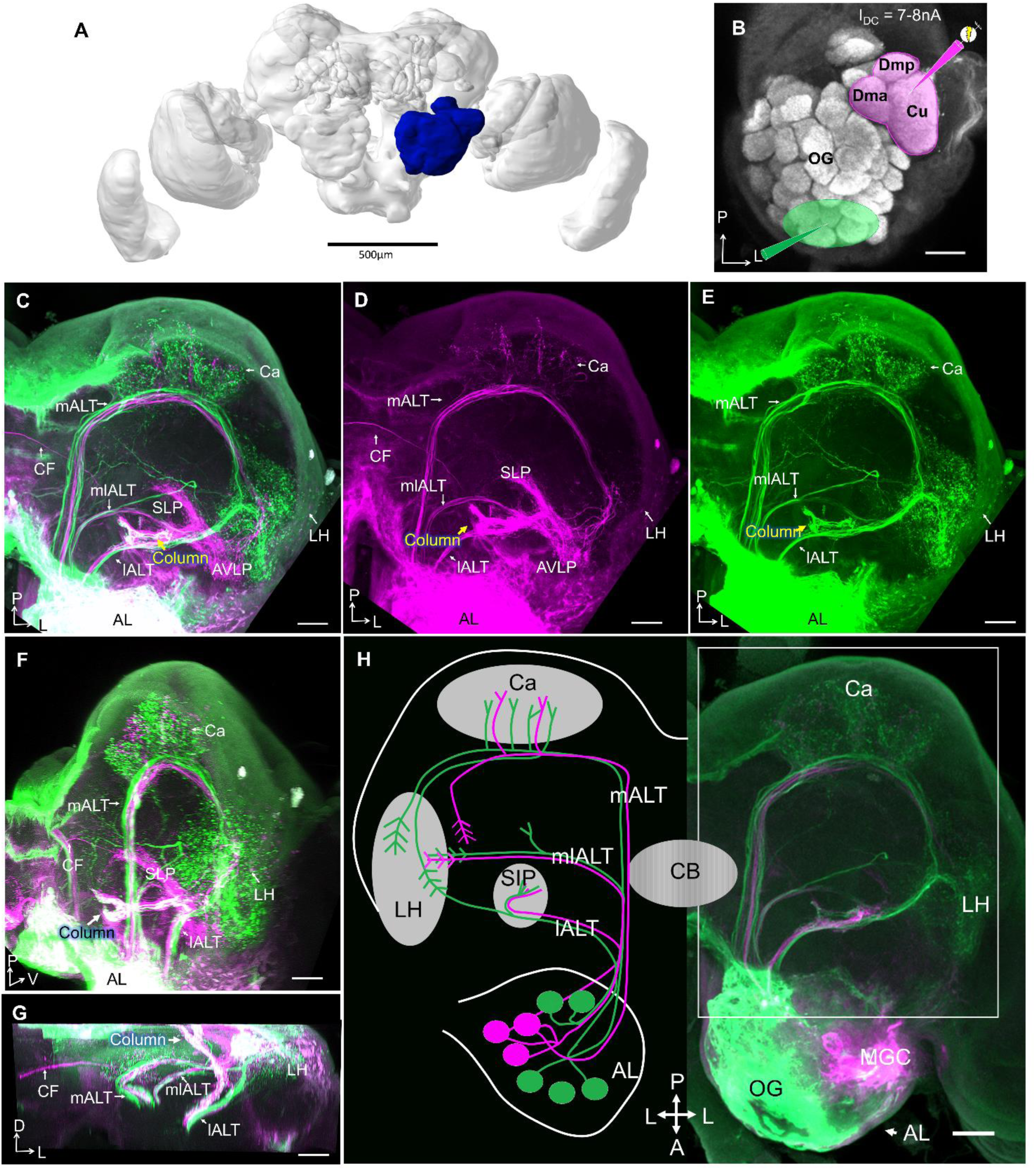
Projection profiles of pheromone and non-pheromone antennal-lobe output neurons in male. **(A)** Brain reconstruction with highlighted left antennal lobe (AL, *blue*). Scale bar, 500 μm. (**B**) Confocal image of the left AL illustrating two ‘anterograde labeling’ sites, the macroglomerular complex (MGC) in *magenta* and the ordinary glomeruli (OG) in *green*. (**C**) Confocal image of a mass-stained preparation showing the three main AL tracts (ALTs), including staining MGC (*magenta*) and the OG (*green*). *White* indicates target regions receiving overlapping input from MGC and OG. Both MGC and OG lateral-tract projection neurons innervate the column (arrow). (**D-E**) Single channel confocal images of (**C**) showing distinct projection patterns of MGC output neurons (**D**) vs. OG output neurons (**E**) in the protocerebrum. (**F-G**) Target areas of lateral ALT (lALT) neurons originating from the MGC (*magenta*) and the OG (*green*) in sagittal (**F**) and dorsal (**G**) view. *White* indicates overlap of projections from MGC and OG. (**H**) Schematic overview of pheromone (*magenta*) and non-pheromone (*green*) AL output neurons in male and a corresponding confocal image. All sections except (**F**) and (**G**) in dorsal view. (l/m/ml)ALT, (lateral/medial/ mediolateral) antennal lobe tract; Ca, Calyces of the mushroom body; LH, lateral horn; PC, protocerebrum; SLP, superior lateral protocerebrum; AVLP, anterior ventro-lateral protocerebrum; CF, commissural fiber. A, anterior; L, lateral; M, medial; P, posterior; V, ventral. Scale bars (**B-H**), 50 μm.

In contrast, the target regions of projection neurons in the lateral ALT overlapped substantially (Fig. 2C). Strikingly, all MGC projection neurons in this tract projected to the column, the small, pillar-like region located in the SIP, tucked between the AOTU and the MB vertical lobe (Fig. 2D). This structure also received projections from lateral-tract projection neurons originating from ordinary glomeruli (Fig. 2E). The column is therefore a site of convergence for pheromone and general odor information. However, only a subset of non-MGC projection neurons in the lateral tract terminate in the column. Approximately half of these cells continue along a lateral trajectory to terminate in the lateral horn, i.e. one of the classic olfactory processing centers (Fig. 2E, H). While the majority of projections remained on the ipsilateral side of the brain, a few commissural fibers originating from the MGC were also observed in these preparations.

Corresponding injections into the female antennal lobe visualized the same ALTs as in males (n = 7), including lateral-tract projection neurons targeting the column and the lateral horn. The commissural fiber bundle found in male (Fig. 2C, D) was not stained in females (Supplementary Fig. 1).

As the column was the only site in the brain that showed substantial overlap in pheromone and plant odor processing neurons, we aimed at obtaining more precise information about the neurons serving as input to this region. We injected a small amount of fluorescent dye to the column region using the intracellular-staining setup (n = 2). These injections resulted in labeling of projection neurons only in the lateral ALT, with no axons marked in the medial or mediolateral ALT (Fig. 3A). This finding confirmed that neurons in the lateral tract provide the sole antennal-lobe input to the column. Furthermore, a belt-like commissural fiber bundle passing posteriorly of the fan-shaped body connected the columns in both hemispheres (Fig. 3B). In addition to the contralateral column, the calyces and the antennal lobe in the contralateral hemisphere were labeled as well (Fig. 3B). In the contralateral antennal lobe, the staining was most pronounced in the cumulus of the MGC, but traces of fluorescent dye were also visible in the two smaller MGC units, as well as in one ordinary glomerulus (Fig. 3C, D). In the preparation with the most numerously labelled neurons, a group of eight cell bodies in the lateral soma cluster of the contralateral antennal lobe were strongly stained (Fig. 3B, red dashed circle), suggesting that at least eight MGC-projection neurons provide input to both columns.

**Figure 3.**
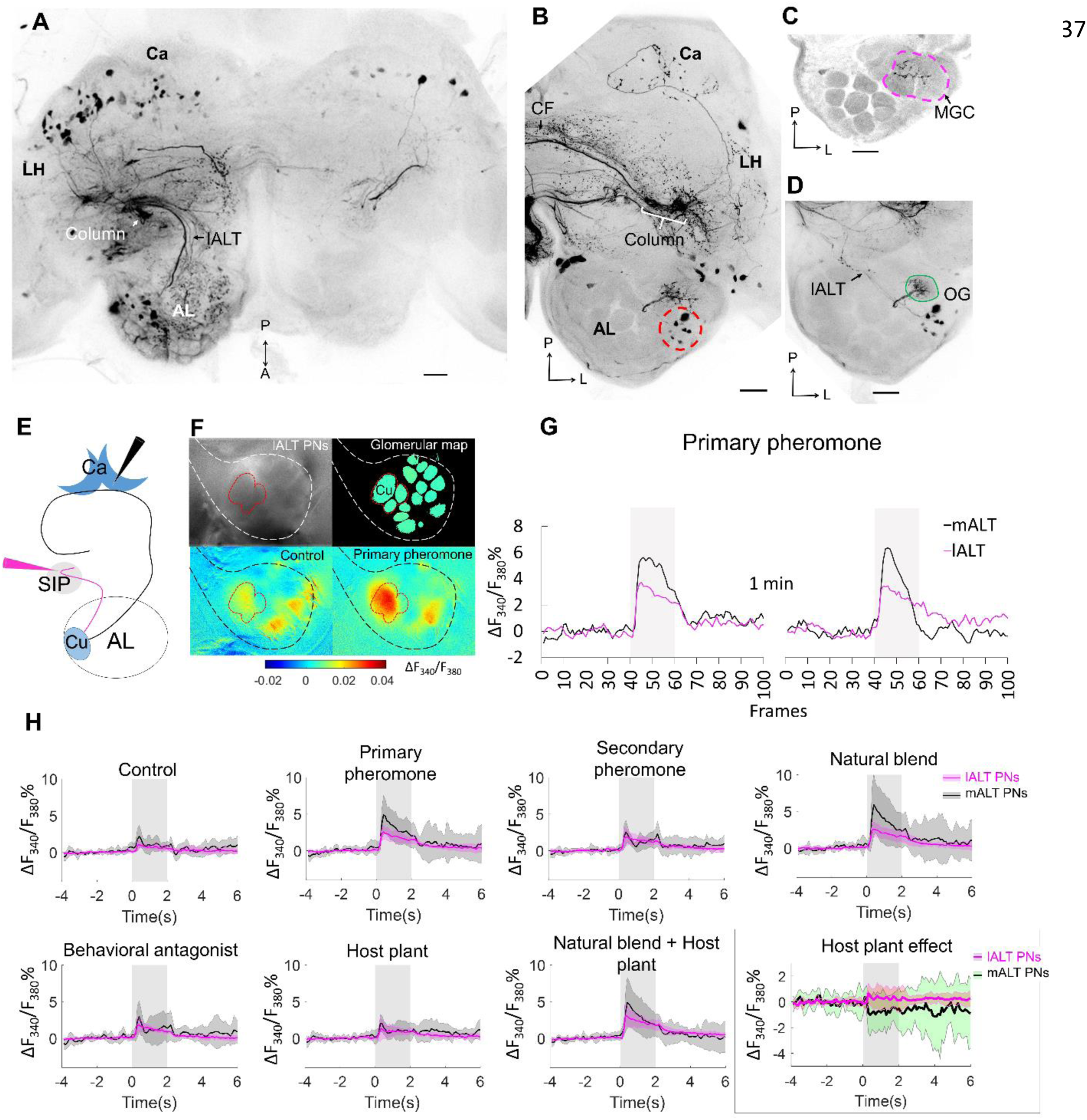
Cumulus neurons confined to the lALT and their odor responses during calcium imaging as compared to mALT neurons. **(A)** Application of dye in the superior intermediate protocerebrum (SIP) visualized antennal-lobe projection neurons confined to the lateral AL-tract (lALT) in the ipsilateral hemisphere. These neurons, which originate from both macroglomerular complex (MGC) and ordinary glomeruli (OG), have somata in the lateral cell cluster and project to the ipsilateral column. (**B**) Dye application in the SIP also labeled a bundle of commissural fibers (CFs) crossing the brain midline and connecting with the contralateral column. Here, in the other hemisphere, eight AL somata (red dash circle) were labeled, demonstrating that these bilateral AL neurons connect the column in both hemispheres. Stained projections in the calyces (Ca) indicate that the bilateral neurons innervate this neuropil as well. (**C-D**) Confocal images showing that the dendrites of the bilateral neurons branch in the MGC (**C**) and one ordinary glomerulus located more ventrally (**D**). (**E**) Illustration of the two ‘retrograde labeling’ sites used for application of the calcium indicator. *Black* arrow head indicates the injection point into the calyx for labeling the mALT neurons. *Magenta* arrow head indicates the injection point in the column region to label the lALT neurons. (**F**) Pictorial material representing calcium imaging data: *top-left*, raw image of an AL stained with Fura from the column; *top-right*, processed image showing a map of recognized glomeruli; *down-left* and *down-right*, Heat maps of responses to the control and primary pheromone. *White* and *black* border circumvents the area of AL, *red* border shows the MGC region. (**G**) Example of calcium imaging traces showing response to the primary pheromone from neurons in the mALT (*black*) and lALT (*magenta*). The standardized traces quantify the neuronal activity of two repeated stimulations with 100 ms sampling frequency. The interval between stimulations is 1 min. (**H**) Responding patterns of lALT and mALT neurons to each stimulus. Host plant effect is illustrated in the *black* rectangle. (l/m)ALT, (lateral/medial) antennal lobe tract; LH, lateral horn; A, anterior; L, lateral; M, medial; P, posterior. Scale bars, 50 μm. *Grey* bar, the duration of the stimulus (2s).

### Neurons projecting to the column functionally differ from those projecting to the calyces

After we established that the column receives exclusive input from the lateral ALT and serves as a likely major site for pheromone processing, we aimed at identifying functional correlates of these anatomical findings. We therefore performed calcium imaging on neurons originating in the main glomerulus of the MGC, the cumulus. We specifically compared the response patterns of cumulus-neuron populations within the lateral tract (projecting to the column) to those of the medial tract (projecting to the calyx). For this purpose, we applied a calcium-sensitive dye (Fura 2) together with Alexa 488 in either one of the two target sites (Fig. 3E). As expected, injection from the calyces selectively labeled medial-tract neurons, while injections from the column region only labeled neurons confined to the lateral tract (Supplementary Fig. 2). The imaging data were obtained from the dorsal region of the antennal lobe, i.e. the input sites for both sets of neurons (Fig. 3F, G).

Cumulus lateral-tract neurons responded to all insect-produced compounds, including the primary pheromone, secondary pheromone, behavioral antagonist, and the natural pheromone blend (Fig. 3H). These responses were similar across all stimulus conditions and showed a consistently tonic temporal profile with a moderate response amplitude. In contrast, the medial-tract neurons projecting to the calyces were activated mainly by stimuli containing the primary pheromone. These responses were stronger and showed a phasic component that decayed over the course of the stimulation period. Interestingly, despite the fact that both MGC neuron populations received input from the same glomerulus, i.e. the cumulus, there were clear differences in response patterns to the pheromone stimuli. In addition, the mixture of plant odors and pheromone blend elicited a suppression in medial-tract neurons, whereas the lateral-tract neurons showed no such effect (Fig. 3H, black rectangle).

### MGC neurons of the lateral ALT are morphologically and physiologically heterogeneous

As we found broad differences in the odor response profiles on the population level between neurons of the lateral and the medial ALTs, we next aimed at investigating how these odor responses were reflected on the level of anatomically identified, single neurons. We thus carried out intracellular recordings from the thick dendrites of MGC projection neurons combined with intracellular dye injection. Ten MGC lateral-tract neurons were identified morphologically and physiologically, all having glomerular dendritic arborizations focused in the cumulus of the MGC. The physiological features of these neurons were then compared to nine MGC cumulus neurons confined to the medial tract.

### Morphological features

All ten MGC lateral-tract projection neurons from which complete dye fills were obtained originated in the cumulus and projected to the column in the SIP. Typically, these neurons exited the postero-ventral part of the antennal lobe and projected laterally. Before reaching the lateral horn, they changed direction and continued dorso-medially, eventually terminating in the column (located between the AOTU and the MB vertical lobe). Despite all targeting the column and having their somata located in the lateral cell body cluster of the antennal lobe, these projection neurons constituted a relatively heterogeneous population. Four were unilateral neurons targeting exclusively the ipsilateral column, whereas the remaining six cells projected bilaterally.

All unilateral neurons turned off their initial course within the lateral ALT and continued dorsally towards the column. Here they formed a mainly unbranched terminal projection (example in Fig. 4A-C; all unilateral neuron anatomies shown in Supplementary Fig. 3A-D). Two of these neurons extended a short side-branch into the lateral protocerebrum before entering the column.

**Figure 4.**
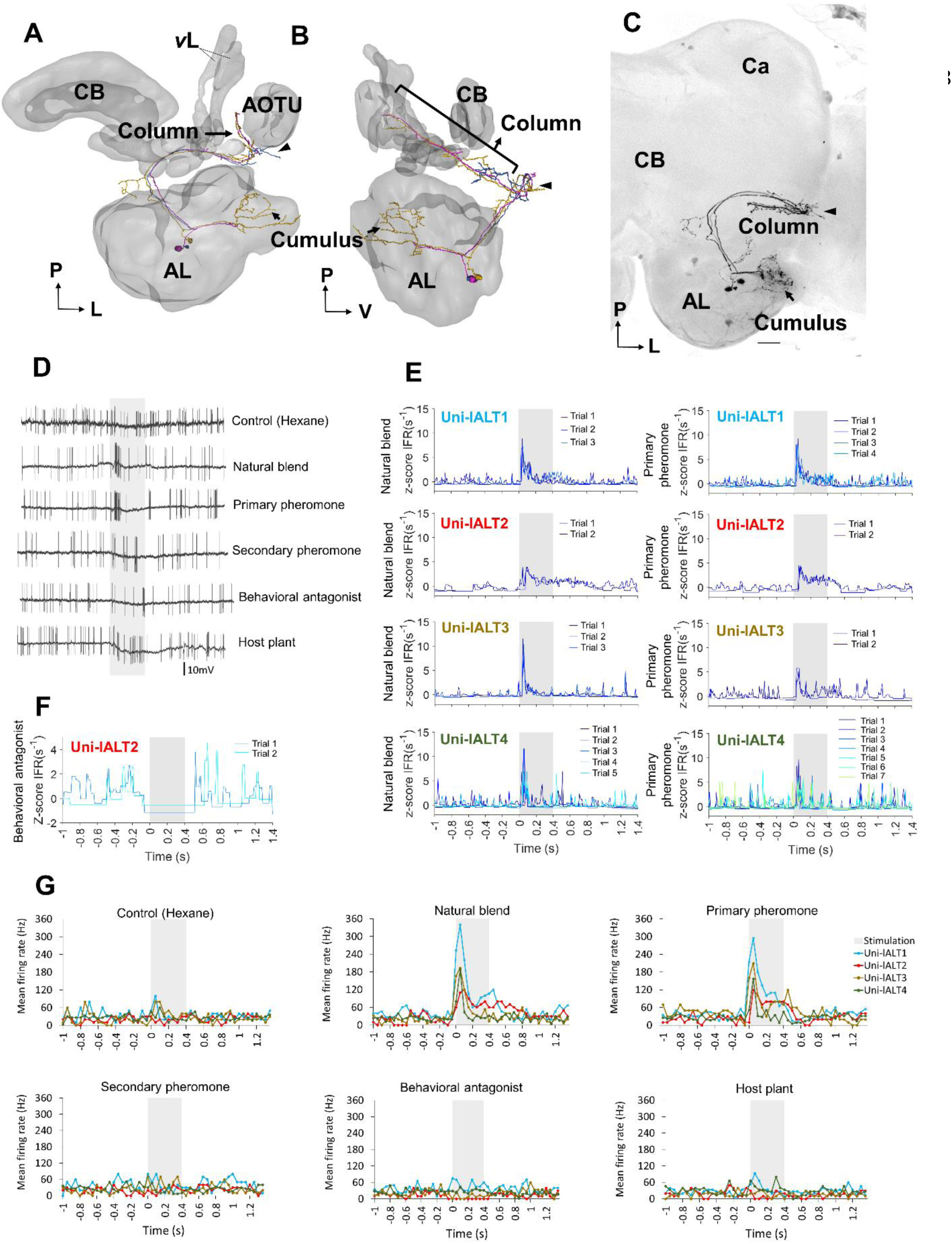
Morphology and electrophysiology of unilateral lateral-tract neurons originating in the cumulus. **(A-B)** 3D reconstruction of three morphologically similar, co-stained neurons (Uni-lALT4) in dorsal-frontal view (**A**) and sagittal view (**B**). The neurons project directly to column tucked between the vertical lobe (*v*L)and the anterior optic tubercle (AOTU). The arrow head indicates a short side branch extending from one axon posterior of the antennal lobe (AL). (**C**) Confocal image of the three lateral-tract cumulus neurons. (**D**) Spiking activity of one of the three unilateral neurons during application of odor stimuli. The neuron showed a phasic activation to the natural blend and the primary pheromone. (**E**) Traces of instantaneous firing rates including responses of four unilateral lALT neurons to the natural blend and the primary pheromone on each trial. (**F**) Traces of instantaneous firing rates of one neuron (Uni-lALT2) showing an inhibitory response to the behavioral antagonist. (**G**) Mean spike frequencies of repeated trials in each neuron (bin-size: 50 ms). Ca, Calyces of the mushroom body; CB, central body. A, anterior; L, lateral; M, medial; P, posterior. Scale bars, 50 μm. *Grey* bar, the duration of the stimulus (400 ms).

The six bilateral neurons targeted the column in *both* hemispheres. Like their unilateral counterparts, they originated in the cumulus and followed the initial course of the lateral ALT. At the turning point adjacent to the lateral horn, the axons split. One projection targeted the ipsilateral column, whereas the other targeted the contralateral column, with the main axon crossing the brain midline posteriorly of the fan-shaped body. In addition to innervating the column in both hemispheres, all bilateral neurons had some terminal branches in the ipsilateral calyx.

Based on their projection patterns, the six bilateral neurons were classified in two sub-populations (all morphologies shown in Supplementary Fig. 4E-J). One consisted of three morphologically similar neurons with clearly defined terminals in restricted areas of the protocerebrum (sub-type I), whereas the other included three morphologically similar neurons with more widespread projections (sub-type II). A typical example of sub-type I is shown in Figure 5. This neuron possesses three main projection fields covering the ipsilateral column, the ipsilateral calyx (Fig. 5C), and the contralateral column. An example of sub-type II is shown in Figure 6. In addition to innervating the three projection fields mentioned above, this neuron had numerous terminal projections in a relatively large area of the ipsilateral protocerebrum and one or two short side branches extending dorsally from the commissural fiber into the SIP, terminating near the medial edge of the MB vertical lobe (Fig. 6D). Finally, this neuron extended numerous short processes from the main axon on its initial route from the antennal lobe.

**Figure 5.**
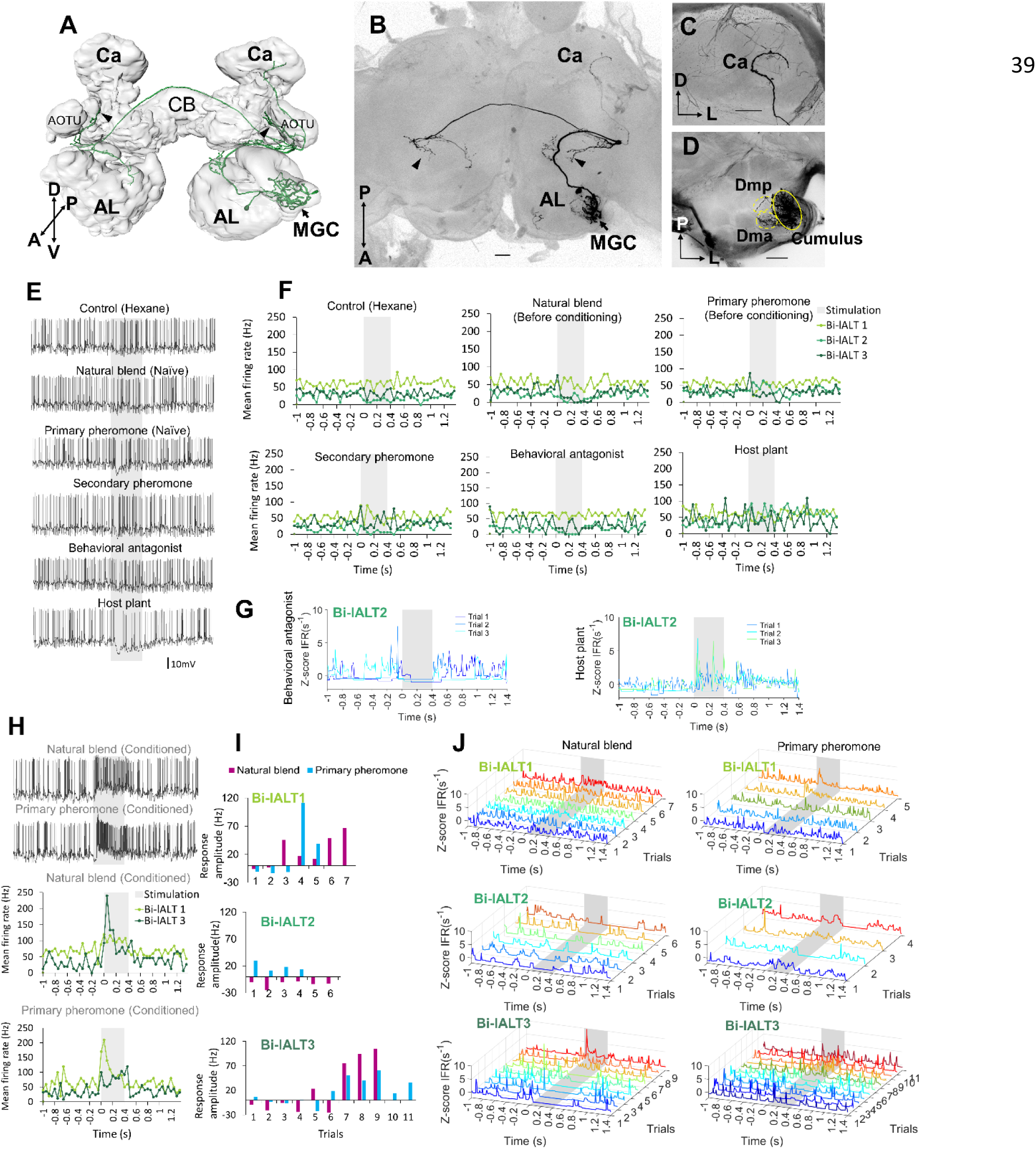
Morphology and electrophysiology of bilateral lateral-tract neurons with restricted projections in the protocerebrum (Sub-type I). **(A)** 3D reconstruction of one neuron (Bi-lALT3) in dorsal-frontal view, demonstrating projections to ipsilateral and contralateral column (arrowheads), and ipsilateral calyx (Ca). (**B-D**) Confocal images of Bi-lALT3. The fiber projecting to the ipsilateral Ca is shown in (**C**). Arborizations in the antennal lobe (AL) are shown in (**D**), including dendrites restricted to the MGC, in which the cumulus is considerably stronger innervated than dma and dmp. (**E**) Spiking activity (bin-size: 50 ms) of one neuron (Bi-lALT3) during first-trial application of each stimulus (i.e., naïve). (**F**) Mean spiking frequencies during repeated trials in the three Sub-type I bilateral neurons before conditioned effect appeared (i.e., first three trials for Bi-lALT1 and first five trials for Bi-lALT3). (**G**) Traces of instantaneous firing rates of one neuron (Bi-lALT2) showing an inhibitory response to the behavioral antagonist and an excitatory response to the host plant. (**H**) Conditioned response in neuron Bi-lALT3. *Upper part:* spiking activities of the neuron during the 7^th^ application of natural blend and primary pheromone, respectively. *Lower part*: spiking frequencies to the same stimuli during 4^th^ -7^th^ trials in neuron Bi-lALT1 and 7^th^ -11^th^ trial in neurons Bi-lALT3. (**I**) Conditioned effect is present in two of the three sub-type I bilateral neurons, i.e. Bi-lALT1 and Bi-lALT3 (no effect in Bi-lALT2). (**J**) Traces of instantaneous firing rates illustrating variable responses to the natural blend and the primary pheromone across repeated trials in the same two Sub-type I bilateral lALT neurons, Bi-lALT1 and Bi-lALT3. AOTU, anterior optic tubercle; CB, central body. A, anterior; L, lateral; M, medial; P, posterior; V, ventral. AL, antennal lobe; Ca, Calyces of the mushroom body; CB, central body. A, anterior; L, lateral; M, medial; P, posterior; V, ventral. Scale bars, 50 μm. *Grey* bar, the duration of the stimulus (400 ms).

**Figure 6.**
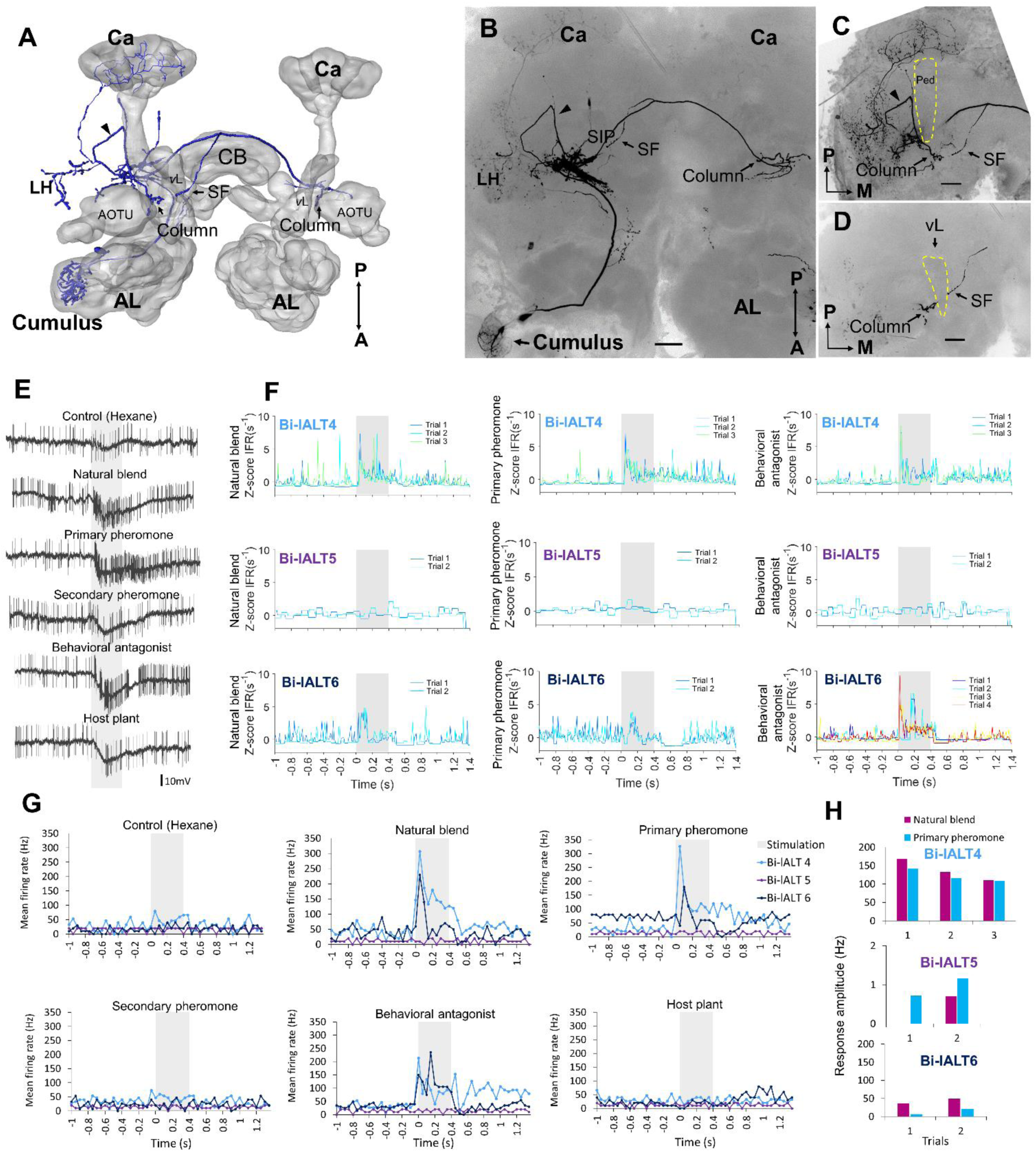
Morphology and electrophysiology of the bilateral lateral-tract neuron type with wide-spread protocerbral innervations (Sub-type II). (**A-D**) 3D reconstruction (**A**) and confocal images (**B-D**) of one neuron (Bi-lALT6) in dorsal-frontal view. In addition to innervating the column in both hemispheres, the neuron projects to calyces (Ca), lateral horn (LH), and superior inferior protocerebrum (SIP) in the ipsilateral hemisphere. The arrow head in panel (**C**) indicates one neural branch forming a small loop close to the lateral edge of the peduncle (Ped) before targeting the ipsilateral column. In panel (**D**), a short side fiber (SF) extending from the commissural fiber and projecting dorsally of the vertical lobe (*v*L) to the ipsilateral SIP is visualized. (**E**) Spiking activity of the bilateral neuron, Bi-lALT6, during application of odor stimuli. The neuron showed a phasic-tonic excitation to the natural blend, the primary pheromone, and the behavioral antagonist. (**F**) Traces of instantaneous firing rates illustrate the responses of the three Sub-type II bilateral lALT neurons to the natural blend, the primary pheromone, and the behavioral antagonist on each trial. (**G**) Mean spike frequencies of repeated trials in each neuron (bin-size: 50 ms). (**H**) No conditioned effects were seen in any of the three Sub-type II bilateral neurons. AL, antennal lobe; CB, central body. A, anterior; L, lateral; M, medial; P, posterior. Scale bars, 50 μm. *Grey* bar, the duration of the stimulus (400 ms).

### Physiological characteristics differ between the two main types of lateral-tract neurons

The two main morphological categories of lateral-tract MGC neurons, i.e. the unilateral and bilateral types, displayed different response patterns during stimulation. Generally, the odor-evoked activation patterns were more homogenous in the unilateral neurons than in the bilateral neurons. All four unilateral neurons displayed significant responses in the form of increased spiking frequencies during stimulation with the primary pheromone and the pheromone mixture (Fig. 4D, E, and G). Three of these neurons (Uni-lALT1, 3, and 4) not only shared consistent responses with respect to odor tuning, but additionally displayed highly similar temporal response profiles. These were characterized by sharp phasic onset responses (lasting 20-30 ms) that gradually faded away towards background activity over the course of the remaining stimulus duration. The fourth neuron (Uni-lALT2) showed a slightly different response pattern. In addition to responding with more phasic-tonic excitation to the primary pheromone and natural blend, it was inhibited by the behavioral antagonist (Fig. 4E, F). Interestingly, the highly phasic responses seen in the majority of unilateral neurons differ from the population responses of lateral-tract neurons as described by calcium imaging. Whereas those experiments revealed a moderate, tonic increase in activation in response to all insect produced stimuli, the single cell responses of the unilateral neurons appeared to be more similar to the calcium-imaging based population responses of neurons in the medial ALT.

In contrast to the relatively consistent response properties of the unilateral MGC neurons in the lateral ALT, the bilateral neurons showed more heterogeneous spiking patterns and less pronounced responses (Figs. 5, 6). The three cells with restricted projections (sub-type I) were not strongly excited by any of the stimuli that elicited responses in the unilateral neurons. Only one neuron, Bi-lALT2, showed an immediate response in the form of a mild excitation to the primary pheromone (and the plant odor) and an inhibition to the behavioral antagonist (Fig. 5G and J). Interestingly, the two remaining cells showed a remarkable switch in response characteristics after repeated stimulations. The first one, Bi-lALT1, displayed no response to the primary pheromone in the first three trials (naïve), but did respond on the forth and the fifth trial (Fig. 5E, H, I, J). A conditioned response occurred during stimulation with the pheromone blend as well, but less pronounced. The second neuron, Bi-lALT3, also displayed conditioned responses, but switched from inhibition to excitation. In this case, the conditioning occurred mainly during repeated stimulations with the natural blend. Here, a phasic-tonic excitation appeared after the sixth trial (Fig. 5I, F). The effect was much weaker during repeated stimulation with the major pheromone alone. The third bilateral neuron with restricted branching pattern, Bi-lALT2, was not dependent on repeated stimulations for eliciting responses. Generally, the responses of these three bilateral neurons had a phasic tonic temporal profile.

The three sub-type II bilateral neurons exhibited even more heterogeneous response patterns than the sub-type I (Fig. 6E). Two of these cells, Bi-lALT4 and Bi-lALT6, showed strong excitatory responses to the primary pheromone and/or pheromone mixture, as well as to the behavioral antagonist, while the third neuron, Bi-lALT5, did not respond to any stimulus. Two of three cells responded immediately, thus repeated stimulations were applied only 2-3 times. They kept a consistent temporal response profile throughout the experiments, characterized by a strong onset burst followed by a lasting tonic response in most stimulus presentations. In some cases, a tonic response (either excitatory or inhibitory) outlasted the stimulus duration. These neurons might therefore account for the tonic responses in the calcium imaging experiments.

Analysis of all neurons’ spontaneous firing properties during the 25-40 s pre-test window revealed a moderately high coefficient of variation of inter-spike interval of the uni-lateral neurons (ISI *C*v =1.14 ± 0.19, for details see supplementary Table 1). This value and the Poisson-like distributions of ISI histograms indicate that the spontaneous firing properties of the unilateral neurons are relatively bursty (Fig. 7 A). This is in contrast to the recorded bilateral neurons, which displayed more varied firing patterns (Fig. 7 B, C). Taken together, the unilateral neurons are homogeneous with regard to both pre-test activity and odor-evoked responses, while the bilateral neurons are heterogeneous according to these properties.

### Lateral-tract and medial-tract neuron physiologies differ in detailed response profiles

So far, our results have shown that the response properties of the lateral-tract neurons differ from those of the medial tract in several ways. On a population level (calcium imaging), responses of the lateral-tract cells occurred during stimulation with all insect-produced compounds tested and were moderate in strength with a largely tonic temporal profile. In contrast, when medial–tract neurons were investigated in the same way, responses were selective to stimuli containing the primary pheromone and showed a more transient temporal profile. On the level of single neurons, the more tonic responses of lateral-tract neurons were partially reflected by individual bilateral neurons, whereas all unilateral neurons of the lateral tract resembled medial-tract neurons more closely. To more thoroughly differentiate the physiological properties of the MGC neurons in the lateral and medial ALT, we quantitatively compared the electrophysiological data from the ten described lateral-tract cumulus neurons with nine medial-tract cumulus-neurons.

A comparison was first conducted between two most homogeneous neuron types, the medial-tract neurons and unilateral lateral-tract neurons. Since the firing pattern of spontaneous activity is an intrinsic property often used to describe individual cell types, we compared the spiking patterns of the two neuron types during the pre-test window. The unilateral neurons in the lateral ALT, in contrast to the medial-tract neurons, displayed more bursty spiking patterns, demonstrated by the shorter minimum ISI during pre-test activity (Mann-Whitney *U* test, *U* = 4, *p* = 0.03), and at the same time, a comparable mean ISIs (*p* > 0.28; Fig. 8A). Moreover, the response amplitude during stimulation with the primary pheromone was larger in the unilateral lateral-tract neurons than in the medial-tract neurons (*U* = 5, *p* = 0.05, Fig. 8B). No other stimuli evoked significantly different firing rate changes between these two neuron types.

**Figure 7.**
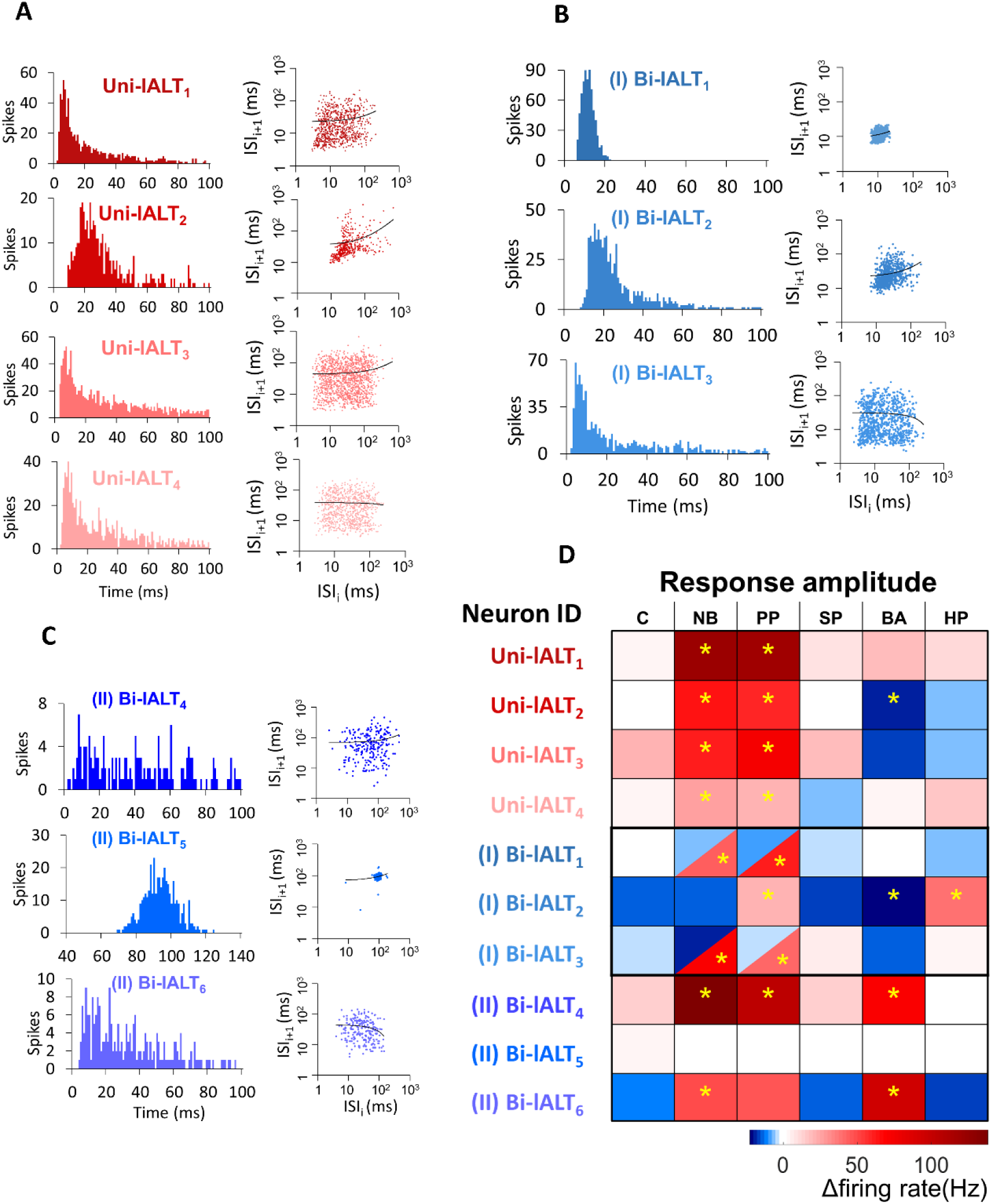
Overview of distinct parameters in all recorded lateral-tract neurons. (**A-C**) Spontaneous firing pattern during pre-test activity, including ISI histograms and Joint ISI scatter plot. (**D**) Heat map of firing rate amplitudes in the 10 lateral-tract neurons. * indicates significant response, determined according to the threshold of baseline activity of individual neurons. C, control; NB, natural blend; PP, primary pheromone; SP, secondary pheromone; BA, behavioral antagonist; HP, host plant.

**Figure 8.**
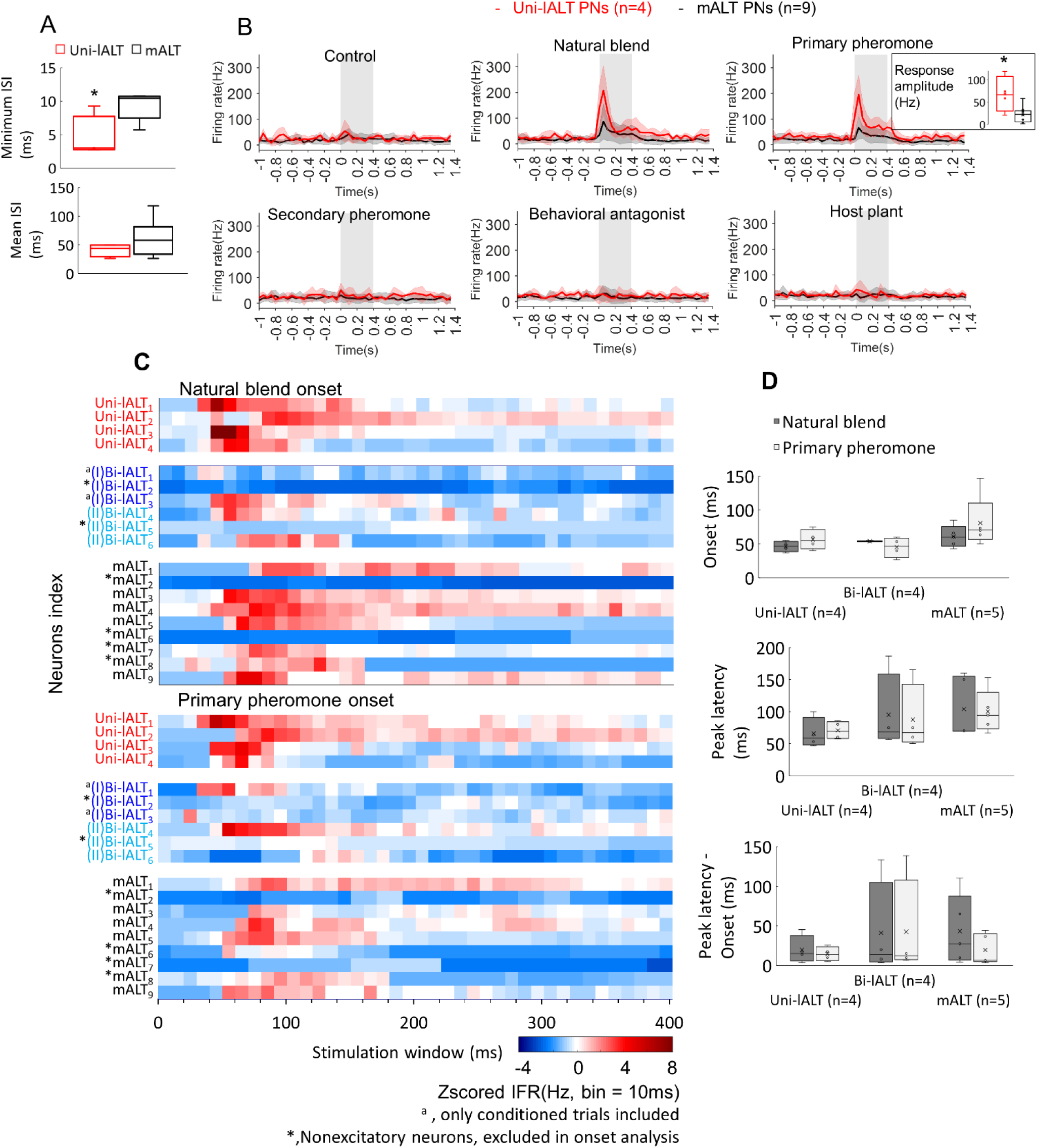
Comparison of spiking activities in lALT neurons and mALT neurons. (**A**) Comparison of minimum and mean ISI in unilateral lALT neurons and mALT neurons before stimulation. (**B**) Average spike frequencies (bin-size: 50 ms) of all unilateral lALT neurons and mALT neurons. *Grey* bar: duration of the stimulus (400ms). The different response strengths of these two types neurons are shown in the inset. (**C**) Heat map of all neurons’ responses during stimulus applications. Each row plots the mean of z-scored IFR (Instantaneous firing rate) of each neuron to the same repeated stimulus during 400 ms stimulation window. (**D**) Latency of response onset and response peak in medial-tract versus lateral-tract neurons with excitatory responses to the natural blend and the primary pheromone.

To compare the odor-evoked responses across all neuron categories, including the bilateral lateral-tract neurons, we compared the instantaneous firing rate plots (binned every 10 ms) of each trial of single-cell recording to all stimuli (Supplementary Fig. 4). Stimuli containing the primary pheromone evoked distinct temporal response patterns in each neuron category, i.e. a phasic response in the unilateral lateral–tract neurons, a phasic-tonic response in the bilateral lateral-tract neurons, and a phasic-tonic or phasic-inhibition response in the medial-tract neurons. The response patterns of unilateral neurons to the main pheromone component and to the pheromone mixture were largely consistent across all four neurons. Medial-tract neurons, on the other hand, responded more variably. Only five of nine medial-tract neurons responded to the primary component, of which four responded to the natural blend. During these stimulations, almost all responsive medial-tract neurons had an apparent longer response delay compared to the lateral-tract neurons (Fig. 8C). To confirm this observation, we quantified these neurons’ onset of excitatory responses based on each recorded trial. (Fig. 8D). The onsets were determined at the time point when the instantaneous firing rate exceeded the response threshold (mean instantaneous firing rate in the 1s pre-stimulation+ 1.96*SD). The response onsets of unilateral neurons (n=4) during stimulation with the primary pheromone were highly consistent and appeared about 30 ms earlier than corresponding onsets in medial-tract neurons (n=5, Fig. 8D). Interestingly, unilateral lateral-tract neurons illustrated a rather short duration between the onset and the peak in comparison of bilateral neurons and the medial tract neurons (Fig. 8D).

Several physiological features, including response profile, temporal response pattern, and onset delay, as well as peak delay, suggest that the unilateral lateral-tract neurons respond more homogeneously than the medial-tract neurons, whereas the bilateral neurons are the most heterogeneous population. This was confirmed by correlating the binned instantaneous firing rate between every two trials of 19 neurons confined to the lateral and the medial tract, respectively (Supplementary Fig. 5). The overall response shapes of the unilateral lateral-tract neurons were, on average, more highly correlated with each other than those of the medial-tract neurons when the primary pheromone and the natural blend were used as stimuli. The overall response shapes of the bilateral lateral-tract neurons, on the other hand, had the lowest mean correlation.

## Discussion

In this study, we combined anatomical and functional experiments to characterize morphological and physiological properties of male-specific projection neurons confined to the lateral ALTs in the moth brain. Different from classic olfactory projections, our data revealed the SIP – a recently defined neuropil flanked by the AOTU and the mushroom body vertical lobe (Ito et al., 2014) – as the main target of MGC neurons projecting along the lateral ALT. More specifically, all neurons terminated in a small area within the SIP called the column. In addition, double mass staining experiments demonstrated that lateral-tract neurons originating from the MGC as well as the ordinary glomeruli provide converging input to the column. The neurons originating in the MGC comprise several morphological types, including unilateral and unusual bilateral neurons. Finally, functional characterization of individual neurons and neuron populations via electrophysiological recordings and calcium imaging demonstrated distinct physiological properties of lateral-tract MGC neurons as compared to corresponding neurons in the medial ALT.

### Major targets of lateral-tract neurons across moth species

Among the moth species most intensely studied, there are obvious differences in lateral-tract projection patterns, in particular regarding the male-specific share. In the silk moth, *B. mori*, the MGC lateral-tract projections target the ‘*delta area’*, a pyramid-shaped region in the inferior lateral protocerebrum (Seki et al., 2005). This triangular area is formed by the male-specific axons confined to the medial and medio-lateral tract as well. Projections originating from the ordinary glomeruli, on the other hand, terminate in the lateral horn, both in males and females. We recently found similar projection patterns of neurons passing along the three main antennal-lobe tracts in the Chinese oak silk moth, *Antheraea pernyi*, (Supplementary Fig. 6A). Notably, these domestic moth species seek the mate by walking or gliding, and are barely capable to fly. In moths performing long-distance flight, however, the lateral tract displays a different projection pattern including a specific target region exclusively for lateral-tract neurons. Our data from both mass staining and individual neuron labelling in male *H. armigera* indicates that all lateral-tract projection neurons from MGC and some from ordinary glomeruli target the column within the SIP. In the classical anatomical study of the ALTs in the tobacco hawk moth, *M. sexta*, Homberg et al. showed that the main sub-category of lateral-tract neurons (named POa) branched off from the lateral path and projected dorsally terminating in the region between the AOTU and MB vertical lobe (1988). The notion that these projections “*seem to branch in several ordinary glomeruli or exclusively in the MGC*” is in full agreement with our findings. Previous studies in heliothine moths have identified “dorsally projecting” lateral-tract projection neurons as well. In a recent study of two heliothine species, *Heliothis virescens* and *Heliothis subflexa*, lateral-tract MGC neurons passing along ‘*the dorsal path*’ were reported (Lee et al., 2018). Besides, similarly projecting lateral-tract neurons that originate from several ordinary glomeruli were found in *H. virescens* (Ian et al., 2016, Rø et al., 2007). In addition, we have found lateral-tract projections targeting the column in the SIP in two migratory moths, the Silver Y, *Autographa gamma*, and the Bogong moth, *Agrotis infusa* (Supplementary Fig. 6B and C). Altogether, the data obtained from flying and non-flying moths indicate that the column in the SIP might be involved in odor-evoked flight behavior, aimed at tracking females or other sources of essential odors over comparably large distances. Interestingly, we noted that *B. mori* and *M. sexta*, i.e. phylogenetically related species belonging to the same superfamily (*Bombyciodea)*, exhibit less similar lateral-tract systems than *M. sexta* and the more distantly related heliothine moths and other noctuids. This indicates that functional requirements rather than phylogenetic distance are the main factors driving the evolution of these projection areas.

A comparable target region of antennal-lobe projections close to the MB vertical lobe was also reported in two other well-studied insect species. In *Drosophila*, the most common lateral-tract neurons (AL-LPN2 in Tanaka et al., 2012) mainly terminate at the ventromedial part of lateral horn and an area surrounding the MB vertical lobe, termed the ‘ring neuropil’. In the honeybee, the mediolateral ALT, i.e. the path likely corresponding to the lateral tract in other insects (Ian et al, 2016), also targets a region called the ring neuropil (Kirschner et al., 2006). While future work will have to show whether all these regions are homologous, it is tempting to speculate that the ring neuropil in the fruit fly and honeybee might be functionally comparable to the column in moths.

### Unilateral lateral-tract neurons could boost robust signal transmission

Based on the homogenous morphology and physiology of the unilateral MGC lateral-tract neurons, we can conclude that they convey highly consistent information to a specific target area. The signals carried by these cells are even more uniform than those of cumulus-neurons confined to the medial tract. In addition, the lateral-tract neurons have shorter response delays. The encoding of a strong, consistent, and fast signal to a narrow target area, as performed by the unilateral neurons, seem unsuitable to capture fine details of the stimulus. In contrast, this configuration might facilitate robust information transmission with a minimum of ambiguity. While medial-tract neurons are involved in odor identification and establishment of odor memory – functions that require fine-tuning and precision - the uni-lateral lateral-tract neurons could serve a more direct role for fast control of key behaviors, requiring strength and sturdiness. Interestingly, these observations are in line with the robustness–efficiency trade-off hypothesis, stating that neural coding cannot be simultaneously optimized for robustness and efficiency, i. e. information capacity (Pryluk et al., 2019). The observed division of labor between neurons of the medial and lateral ALT thus might be an implementation of this tradeoff.

### Bilateral MGC lateral-tract neurons are suited to optimize pheromone tracking strategies

A significant proportion of the lateral-tract MGC projection neurons stained in this study (six of ten) were bilateral neurons targeting the column in both brain hemispheres. When adding dye to the column, eight MGC-connected somata in the lateral cell cluster of the contralateral antennal-lobe were labeled (Fig. 3B), indicating the presence of at least eight bilateral projection neurons confined to the lateral tract. Considering that there are approximately 32 lateral-tract MGC neurons innervating the column in *M. sexta* (Homberg et al. 1988), and assuming equivalent cell numbers in the heliothine moth, the bilateral type may constitute approximately 25% of the total number of MGC lateral-tract neurons in *H. armigera*.

The bilateral lateral-tract projection neurons identified in *H. armigera* seem to be present exclusively in males. As male-specific lateral-tract neurons with bilateral innervations in the SIP region were also found in three other moth species, *H. virescens* (Ian et al., 2016), *Heliothis subflexa* (Lee et al., 2018), and *Agrotis segetum* (Wu et al., 1996), this neuron category likely plays a role specific to pheromone processing. One putative role is to integrate pheromone input with bilateral signals from other sensory modalities. Male moths seek the calling female not by sensing a chemical gradient of pheromone concentration, but by combining vision and mechano-sensation to fly against the wind as long as the pheromone signal is present. Both the visual and mechano-sensory systems are inseparably linked to space and thus to bilateral coding mechanisms (Pfeiffer and Homberg, 2014, Homberg et al., 2011, Patella and Wilson, 2018, Jacobs et al., 2008). An alternative explanation is that the bilateral projections summate simultaneous bilateral signals (Rodrigues, 1988) and thereby double the input from upstream sensory neurons to enhance signal-to-noise ratio (Raman et al., 2008).

Interestingly, among the bilateral neurons identified here, two of sub-type I enhanced their responses during repeated exposure to pheromone stimuli. These MGC bilateral projections could therefore serve as an arousal system that maintains plume tracking behavior by switching the downstream circuit into a more responsive state once the signal has crossed a reliability threshold. This strategy would prevent continued tracking after the detection of a single pheromone pulse, averting expensive investment into the pursuit of potentially false-positive signals. Only when several odor filaments are detected in close sequence, the neurons would fire and switch the circuit to drive continued seeking of the odor source. Consistent with this hypothesis, behavioral tests in a wind tunnel showed that when a single pheromone pulse was applied, male *H. virescens* followed a shorter distance than when multiple pulses of pheromone were used as stimulus (Vickers and Baker, 1994).

### A potential role of the column in odor integration and navigation

Both MGC and non-MGC lateral-tract projection neurons target the column. Whereas the MGC projection neurons are mainly uni-glomerular, those responding to plant odors have been described as multi-glomerular (Ian et al., 2016, Rø et al., 2007, Homberg et al., 1988). A multi-glomerular neuron will only be fully activated when the stimulus contains all individual components corresponding to the combination of the innervated glomeruli (Løfaldli et al., 2012). Thus, this arrangement suggests that all lateral-tract neurons projecting to the column are “pre-coded” in the antennal lobe to transmit specific, behaviorally significant information about pheromones, distinct plant-odor combinations, or a combination of both. Which information besides pheromones is encoded via this labelled line pathway remains to be shown.

One behavior in which such labeled lines might be of importance is odor-evoked spatial orientation, e.g. to follow a conspecific pheromone plume, to avoid detrimental odors, or to pursue egg-laying sites. While not being directly connected (Supplementary Fig. 7), the spatial proximity of the column to visual processing neuropils (e.g. the AOTU) is intriguing, as both might share downstream targets. Flying insects rely fundamentally on the visual system when tracing an odor source (reviewed by Baker and Hansson, 2016). Without visual feedback, moths find it very difficult to track airborne pheromone plumes successfully (Willis et al., 2011). Contrary to intuition, no odor-mediated behavior in moths is based on chemotaxis, i.e. there is no direct navigating response to odor concentration gradients. Moths rather steer against the odor source in correspondence with optomotor anemotaxis (anemo: wind, taxis: directed movement), which requires both olfactory and visual feedback (Kennedy and Marsh, 1974). Lateral-tract projection neurons targeting the column may therefore provide olfactory signals optimized for being integrated with visual information and might therefore be directly involved in triggering species-specific, odor-evoked upwind flight behavior. To explore the putative integration of visual and olfactory input in neural networks connected to the column provides an exciting subject for future studies.

## Materials and Methods

### Insects

Male and female moths (2-3 days) of *H. armigera* (Lepidoptera: Noctuidae; Heliothinae) were used in this study. Pupae were supplied by Keyun Bio-pesticides (Henan, China). After emergence, the moths were kept at 25 °C and 67% humidity on a 14:10 h light/dark cycle (lights on at 18:00), with 10% sucrose solution available ad libitum. According to Norwegian law of animal welfare, there are no restrictions regarding experimental use of Lepidoptera.

### Intracellular recording and staining

Preparation of the insect has been described in detail elsewhere (see Rø et al., 2007, Zhao et al., 2014). Briefly, the moth was restrained inside a plastic tube with the head exposed and then immobilized with dental wax (Kerr Corporation, Romulus, MI, USA). The brain was exposed by opening the head capsule and removing the muscle tissue. The procedure of intracellular recording/staining of antennal-lobe projection neurons was performed as previously described (Zhao et al., 2014, Ian et al., 2016). Sharp glass electrodes were made by pulling borosilicate glass capillaries (OD: 1 mm, ID: 0.5 mm, with filament 0.13 mm; Hilgenberg GmbH, Germany) on a horizontal puller (P97; Sutter Instruments, Novarto, CA, USA). The tip of the micro-pipette was filled with a fluorescent dye, i.e. 4% biotinylated dextran-conjugated tetramethylrhodamine (3000 mw, micro-ruby, Molecular Probes) in 0.2 M potassium acetate (KAc). The glass capillary was back-filled with 0.2 M KAc. To facilitate microelectrode insertion into the tissue, the sheath of the antennal lobe was gently removed by using fine forceps. The exposed brain was continuously supplied with Ringer’s solution (in mM: 150 NaCl, 3 CaCl2, 3 KCl, 25 sucrose, and 10 N-tris (hydroxymethyl)-methyl-2-amino-ethanesulfonic acid, pH 6.9). A chloridized silver wire inserted into the muscle in the mouthpart served as reference electrode. The recording electrode, having a resistance of 70–150 MΩ, was carefully inserted into the dorsolateral region of the AL via a micromanipulator (Leica). Neuronal spike activity was amplified (AxoClamp 2B, Axon Instruments, Union, CA) and monitored continuously by oscilloscope and loudspeaker. Data were digitized with CED1401 micro using Spike2 6.02 (Cambridge Electronic Design, Cambridge, England) as acquisition software. During recording, the moth was ventilated constantly with a steady stream of fresh air. Spontaneous activity in the pre-test window was recorded for a period of 25–40 seconds. During odor stimulation, a pulse of air from the continuous airstream was diverted via a solenoid-activated valve (General Valve Corp.) through a glass cartridge bearing the odorant on a piece of filter paper. Six odors were tested in each recording experiment. The stimulation period was 400 ms, and application of each odor was repeated at least two times. After testing all odor stimuli, the neuron was iontophoretically stained by applying 2–3 nA pulses with 200 ms duration at 1 Hz for about 5–10 min. In order to allow neuronal transportation of the dye, the preparation was kept overnight at 4 °C. The brain was then dissected from the head capsule and fixed in 4% paraformaldehyde for 1 h at room temperature before it was dehydrated in an ascending ethanol series (50%, 70%, 90%, 96%, 2 × 100%; 10 min each). Finally, the brain was cleared and mounted in methyl salicylate.

### Mass staining

Three types of mass staining experiments were performed. In the first, we achieved an overview of pheromone projection neurons versus non-pheromone projection neurons in male moths via double dye injection experiments. Two fluorescent dyes were applied to the same preparation – one to the MGC and the other to the ordinary glomeruli. A focal injection was achieved by inserting a sharp electrode filled with 4% micro-ruby solution to the MGC region, using our intracellular electrophysiology setup and applying depolarizing current pulses of 7-8 nA at 1 Hz for 15 min. To label non-MGC projection neurons, the antennal lobe region including ordinary glomeruli was manually perforated with a fine needle containing crystals of Alexa Fluor 488 dextran (10000 mw, Molecular Probes). To visualize input and output areas of the male column region, micro-ruby was applied to this neuropil in the second experiment via pulsed current injection. To compare the antennal-lobe output projections in male and female, a third mass staining experiment was conducted by applying micro-ruby to the female antennal lobe. In all mass staining experiments, the brains were kept for 2 h at room temperature for transportation of the dyes. The subsequent procedure included dissection, fixation, dehydration, and mounting in methyl salicylate as described above.

### Calcium imaging

Totally, 16 males (age: 2-3 days) were used to measure pheromone-evoked responses from the MGC by means of calcium imaging. Selective staining of projection neurons in *H. armigera* was described elsewhere (Ian et al., 2017). Briefly, the head capsule was opened after the moth was immobilized in a small plastic tube. Membranes and trachea covering the brain were gently removed. Glass electrodes loaded with calcium-sensitive dye, Fura-2 dextran (potassium salt, 10000 mw, Molecular Probes) was inserted into the column in eight of the individuals and into the calyces in the eight remaining, to stain (via retrograde transport) lateral-tract neurons and medial-tract neurons, respectively. In order to determine whether the retrograde staining was successful, Fura-2 was mixed with a fluorescent dye, Alexa 488 dextran, having the same molecular weight as the calcium indicator. Then the insect was kept in the dark at 4°C overnight.

*In vivo* calcium imaging recordings were obtained from the dorsal region of the antennal lobe with an epifluorescent microscope (Olympus BX51WI) equipped with 20x/1.00 water immersion objective (OlympusXLUMPlanFLN). Images were acquired by a 1344x1224 pixel CMOS camera (Hamamatsu ORCA-Flash4.0 V2 C11440-22CU). The preparation was excited with monochromatic light of 340 nm and 380 nm, respectively (TILL Photonics Polychrome V). Data were acquired ratio-metrically. A dichroic mirror (420 nm) and an emission filter (490-530 nm) were used to separate the excitation and emission light. Each recording consisted of 100 double frames at a sampling frequency of 10 Hz, with 43 ms and 14 ms exposure times for the 340 nm and 380 nm lights, respectively. The duration of one recording trial was 10 s, including 4 s with spontaneous activity, 2 s odor stimulation, and a 4 s post-stimulus period. The odor stimulation was carried out by a stimulus controller (SYNTECH CS-55), via which humidified charcoal filtered air was delivered through a 150 mm glass Pasteur-pipette with the stimulus on a piece of filter paper inside. Each odor stimulus was applied twice. To avoid possible adaptation the interval between trials was 60 s. To confirm that the obtained calcium imaging corresponded to neurons confined to the medial and lateral tract, respectively, we dissected 50% of the brains afterwards to visualize the traces of Alexa 488 dextran.

### Odor stimulation

During intracellular recordings, the following stimuli were tested: (i) the primary sex pheromone of *H. armigera*, Z11-16:Al, (ii) the secondary sex pheromone, Z9-16:Al, (iii) the binary mixture of Z11-16:Al and Z9-16:Al, (iv) the behavioral antagonist of *H. armigera*, Z9-14:Al, (v) the head space of a host plant (sunflower leaves), and (vi) hexane as a vehicle control. The three insect-produced components were obtained from Pherobank, Wijk bij Duurstede, Netherlands. The mixture of Z11-16:Al and Z9-16:Al were in a 95:5 proportion to resemble the natural blend emitted by conspecific females (Kehat et al., 1980, Piccardi et al., 1977, Wu et al., 1997). Stimuli i-iv were diluted in 99% hexane (Sigma) with a final concentration of 500 ng/ml. Twenty μl of each stimulus were applied to a filter paper placed inside a 120 mm glass cartridge. For the female-produced compounds, this meant that each filter paper contained 10 ng of the relevant stimulus. The same odor stimuli as listed above were used during the calcium imaging experiment, but at a higher concentration (required to evoke a response in this technique), i.e. 10 µg at the filter paper. An additional stimulus containing a 50:50 mixture of host plant (20µl) and pheromone mix (20µl) was added in the calcium imaging measurements.

### Immunohistochemistry

A total number of nine brains were synapsin-stained in order to generate representative data of the brain of the *H. armigera* male. The moth brain was dissected and immediately transferred into a Zinc-Formaldehyde fixative (Ott, 2008) at room temperature overnight. The brain was then washed in HEPES-buffered saline (HBS, 8x30 min) (Ott, 2008), and subjected to a permeabilization step (60 min incubation with a fresh mixture of 20% DMSO and 80% methanol) before being washed 3 x 10 min in Tris-HCL buffer (0.1M, pH 7.4). After pre-incubation in 5% normal goat serum (NGS, Sigma St. Louis, MO, USA) in 0.1 M phosphate-buffered saline (PBS, pH 7.2) containing 0.3% Triton X-100 (PBT), the brain was incubated for 5-6 days at 4°C in the primary antibody, SYNORF1 (dilution 1:25 in PBT containing 1% NGS). Following rinsing in PBT 8x30 min, the brain was incubated for 4-5 days at 4°C with Alexa Flour plus 647 conjugated goat-anti-mouse secondary antibody solution (Invitrogen, Eugene, OR; dilution 1:300 in PBT with 1% NGS). After washing 4 x 30 in PBT and 2 x 30 min in PBS, the brain was dehydrated in increasing ethanol series (50%, 70%, 90%, 95% and 100% (2x), 15 min each). Then, the brain was transferred to the mixture of methyl salicylate and ethanol (1:1) for 15 min and after that cleared completely in methyl salicylate for at least one hour. Finally, the brains were mounted in Permount between two coverslips, separated by spacers.

### Confocal microscopy

Whole brains containing injected neurons were imaged dorso-frontally by using a confocal laser scanning microscope (LSM 800 Zeiss, Jena, Germany) equipped with a Plan-Neofluar 20x/0.5 objective. Micro-ruby staining was excited with a HeNe laser at 553 nm and the fluorescent emission passed through a 560 nm long-pass filter. Staining of Alexa Fluor 488 was excited with an argon laser at 493 nm and a 505-550 nm band pass filter. In addition to the fluorescent dyes, the auto-fluorescence of endogenous fluorophores in the neural tissue was imaged to visualize relevant structures in the brain containing the stained neurons. Since many auto-fluorescent molecules in the tissue are excited at 493 nm, images were obtained using the 493 nm argon laser in combination with a 505-550 nm band pass filter. Serial optical sections with resolution of 1024 x 1024 pixels were obtained at 2 µm intervals through the entire depth of brain. The confocal images shown in this study were edited in ZEN 2.3 (blue edition, Carl Zeiss Microscopy GmbH, Jana, Germany).

The immunostained brains were imaged using a Leica SP8 confocal microscope equipped with a 20x multi-immersion objective (HC PL APO CS2 20x/0.75 IMM). The samples were excited with a 638 nm laser. Images (8 bit) were obtained via the Hybrid detector (standard mode) at a voxel size of 0.76 x 0.76 x 1 µm^3^. Multiple Scans were carried out from both anterior and posterior sides to cover the entire brain.

### 3D brain reconstruction and single neuron tracing

The criteria for selecting suitable preparations for 3D reconstruction were the least damage in regions of interest and sufficient contrast through the entire stack. Out of totally nine immuno-stained male brains, three were selected to construct either full brains or all neuropils of the central brain. Raw confocal stacks were aligned and stitched together using FIJI (Preibisch et al., 2009) and resampled to 1.5x1.5x1.5 µm voxel size in Amira (Amira 5.3; Thermo Fisher, Visualization Science Group). These complete image stacks were then utilized to carry out image segmentation in Amira. We manually labeled key cross sections of each neuropil of interest in all three spatial planes and then used the wrap-tool in Amira to obtain full neuropil volumes. This process yielded segmented image stacks containing all major neuropils of the moth brain. A surface model of the segmented image stacks was produced in Amira. These were either visualized in Amira or exported as obj-files and uploaded to the InsectBrainDatabase (www.insectbraindb.org) for visualization.

Five stained neurons from three preparations were traced manually using the *SkeletonTree* plugin in Amira (Schmitt et al., 2004, Evers et al., 2005). Based on background fluorescence of the neuron channel, neuropils close to the traced neurons were segmented in each brain preparations as described above.

### Nomenclature

For naming the neuropil structures of the brain, we used the nomenclature established by Ito et al. (2014). However, with respect to the lateral horn (LH), we have restricted this region to include the area targeted by the non-MGC uni-glomerular projection neurons passing in the medial ALT. The definition of the LH as the target region of all antennal-lobe projection neurons, as stated in Ito et al. (2014), is not applicable to moths, as a prominent branch of the lateral ALT projects to a region located in the SIP. For naming the sub-classes of projection neurons confined to the various ALTs, we have used the system established by Homberg et al. (1988), however, adjusted to the new names of the tracts. The orientation of all brain structures is indicated relative to the body axis of the insect, as in Homberg et al. (1988).

### Spike data analysis

The electrophysiological data were analyzed in Spike 2.8. When stable neuronal contact was established, we measured the pre-test activity for 25-40 s to determine the basic firing properties of each recorded neuron before applying stimuli. The spike-trains were abstracted from voltage waveform traces when there was a good signal-to-noise ratio which fitted to a single wave-mark. The pre-test activity pattern of each neuron was described by mean interspike interval (ISI), mean firing rate, coefficient of variation (*Cv*) of ISI, ISI distribution, and joint-ISI scatter plot (a time line indicating the relevant intervals during each recording is shown in Fig. 8 in Supplementary material.). Each odor application trial comprised a total period of 2.4 s, including 1 s baseline activity prior the stimulus onset, 0.4 s stimulation period, and 1 s post-stimulation period. For describing neural activity during repeated trials of the same stimulus, mean odor traces showing the neuron’s firing rates every 50 ms were generated. In order to characterize the temporal responding pattern, we also calculated the binned instantaneous firing rate (BIFR) of every 10 ms for each trial.

To measure responses of individual projection neurons, the odor-evoked response properties were analyzed in two steps. (i) For determining significant responses of individual trials, an upper and lower threshold was calculated according to the mean BIFR in the 1 s pre-stimulus (1 s baseline activity) window (BIFRPS):

Upper Threshold_binned instantaneous firing rate_ = mean of BIFRPS + standard deviation of BIFRPS * 1.96

Lower Threshold_binned instantaneous firing rate_ = mean of BIFRPS - standard deviation of BIFRPS * 1.96

If there was an individual BIFR in the stimulation window higher than the value of ‘Upper Threshold’ or lower than ‘Lower Threshold’, the trial was determined as an excitatory or inhibitory response, respectively. If the ‘Lower Threshold’ was less than zero, zero was used as the ‘Lower Threshold’.

(ii) For determining significant responses of repetitive trials, we calculated an upper and lower threshold around the mean firing rate in the pre-stimulus window (FRPS) for each mean odor histogram when there was at least one trial with excitatory/inhibitory response:

Upper Threshold _mean odor histogram_ = mean of FRPS + standard deviation of FRPS * 1.96

Lower Threshold _mean odor histogram_ = mean of FRPS - standard deviation of FRPS * 1.96

When the mean firing rate in the stimulation window was higher than the value of ‘Upper Threshold’ or lower (equal included) than ‘Lower Threshold’, the neuron was determined to display a significant excitatory or inhibitory response, respectively. If the ‘Lower Threshold’ was less than zero, zero was used as the ‘Lower Threshold’.

For displaying the response amplitude, we first standardized the baseline activity by setting the firing rate before stimulation onset to zero. Then, we calculated the spike frequency, i.e. Δ firing rate, averaged over the 400 ms stimulation window. For comparing the odor-evoked response to the same stimulus across individual trials, pairwise Pearson correlations were conducted on the binned instantaneous firing rate histograms of every two trials. The ISI, the firing rate of individual neurons, and the response onset all failed to follow a normal distribution, thus a nonparametric analysis (Mann-Whitney *U* test) was conducted among medial-tract and lateral-tract neurons. All probabilities given are two-tailed. For statistical analysis, SPSS, version 24, was used.

### Calcium imaging data analysis

In this study, neural activities of projection neurons innervating the cumulus of the MGC were analyzed. Recordings were acquired with Live Acquisition V2.3.0.18 (TILL Photonics) and imported in KNIME Analytics Platform 2.12.2 (KNIME GmbH, Konstanz, Germany). Here, ImageBee neuro plugin (Strauch et al., 2013) was used to construct AL maps and glomerular time series. To determine an average baseline activity, the Fura signal representing the ratio between 340 and 380 nm excitation light (F340/F380) from 0.5 to 0.25 s (frames 5-25, within 4 s spontaneous activity) was selected and set to zero. Responses were defined as changes in fluorescent level being significantly different from that corresponding to the average baseline activity, specified as ΔF340/F380.

## Acknowledgements

We thank Jonas H. Kymre (NTNU), Christoffer Nerland Berge (NTNU), and Pramod KC (NTNU) for assistance with data collection. The authors are grateful for financial support from the following organizations: the Norwegian Research Council (project nr. 287052 to B.G.B.), the Swedish Research Council (Vetenskapsrådet; 621-2012-2213 to S.H.), and the European Research Council (ERC) under the European Union’s Horizon 2020 research and innovation program (grant agreement no. 714599 to S.H.).

## Author contribution

XC: Conceptualization, Data collection, Formal analysis, Investigation, Visualization, Methodology, Writing—original draft, Writing—review and editing

SH: Formal analysis, Investigation, Visualization, Supervision, Writing—review and editing EI: Investigation, Visualization

BGB: Conceptualization, Formal analysis, Investigation, Supervision, Funding acquisition, Writing—original draft, Writing—review and editing

## Supplementary material

**SM Fig.1.**
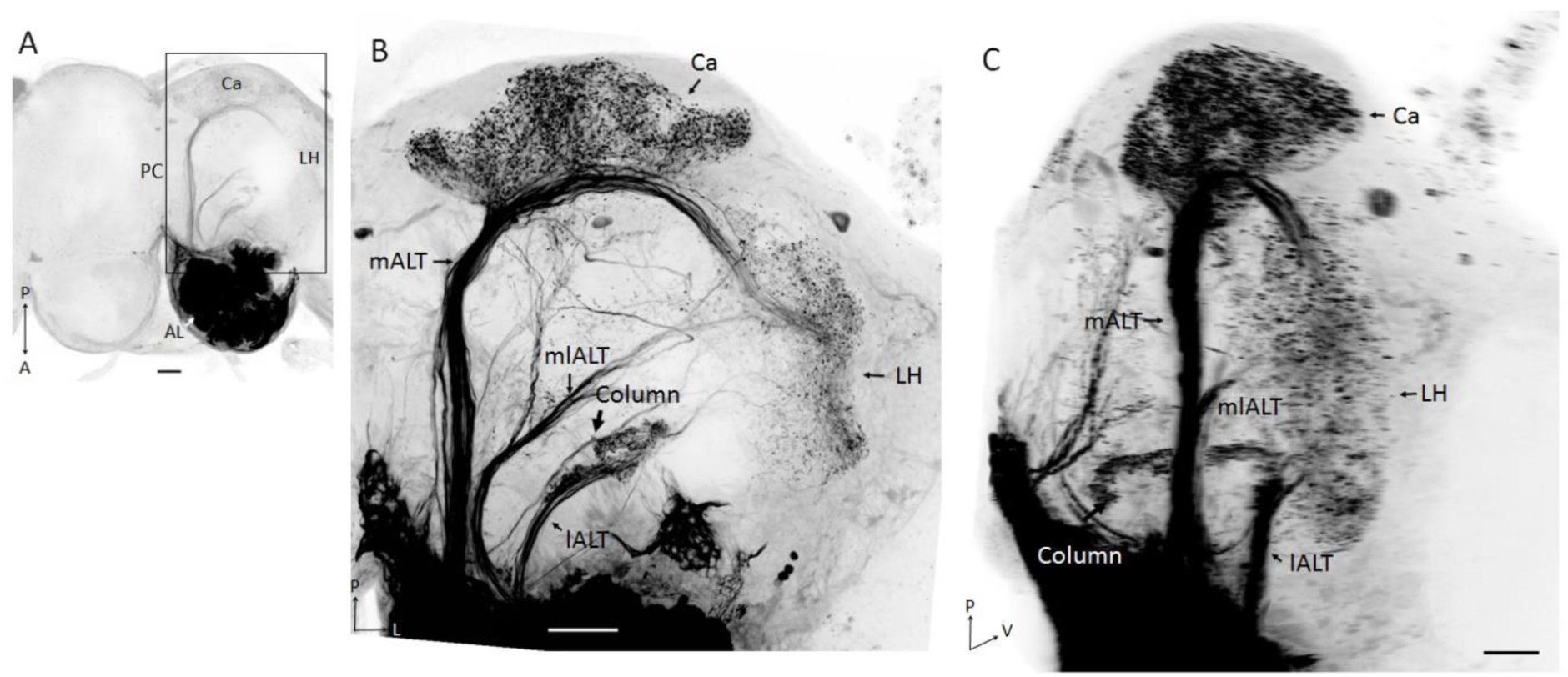
Projection profiles of antennal lobe. (AL) output neurons in female. (**A**) Confocal image of the left AL illustrating the ‘anterograde labeling’ sites in the AL. (**B-C**) Confocal images of the mass-stained preparation showing the three main AL tracts (ALTs) in dorsal view (**B**) and sagittal view (**C**). The strongly labeled column indicates that this area is a main target of lateral-tract neurons in the female moth. However, no commissural fiber bundle originating from the AL was visible here. (l/m/ml)ALT, (lateral/medial/ mediolateral) antennal lobe tract; Ca, Calyces of the mushroom body; LH, lateral horn; A, anterior; L, lateral; M, medial; P, posterior; V, ventral. Scale bars, 50 μm.

**SM Fig.2.**
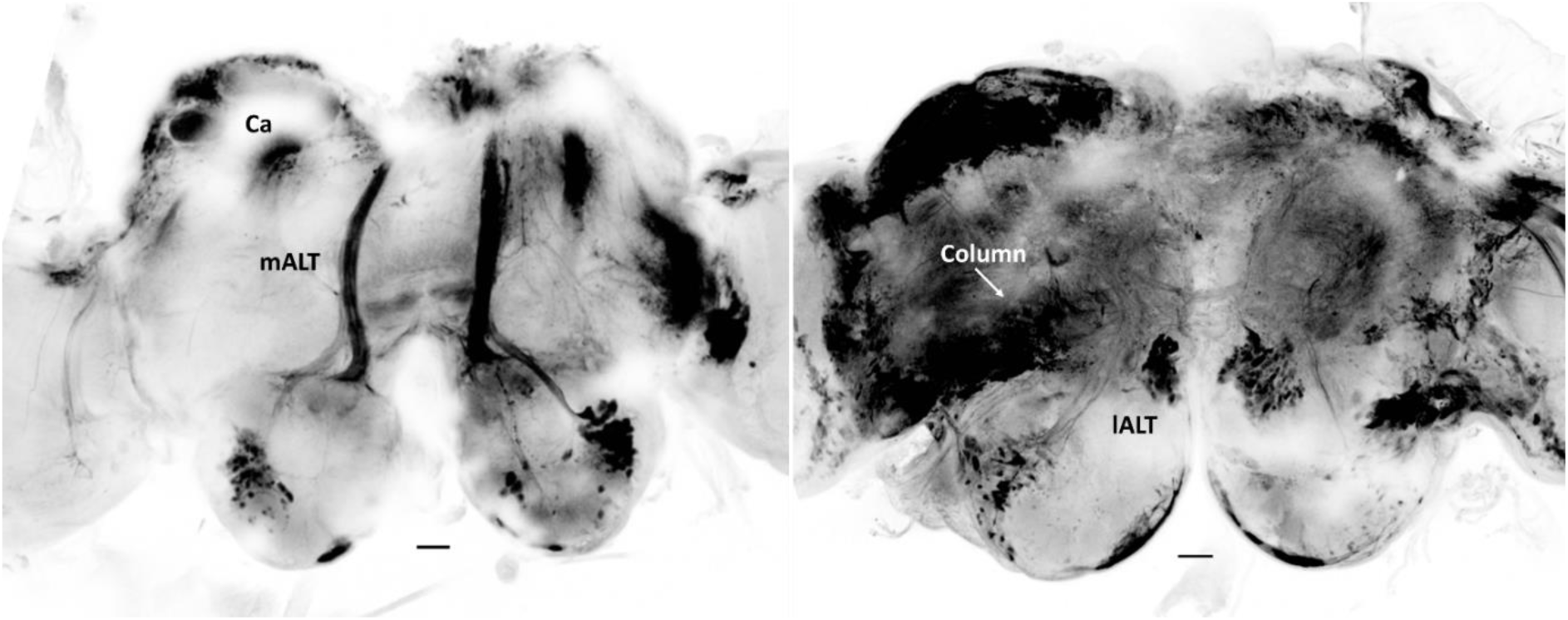
**Selective retrograde labeling of projection neurons in preparations used for calcium imaging experiment (dorsal view). *Left***: exclusive labeling of medial-tract neurons when the mixture of Fura-2 and AF488 was applied into the calyces (Ca). ***Right***: labeling of lateral-tract neurons when we the dye mixture was applied into the superior intermediate protocerebrum (SIP). (l/m)ALT, (lateral/medial) antennal lobe tract; Scale bars, 50 μm.

**SM Fig. 3.**
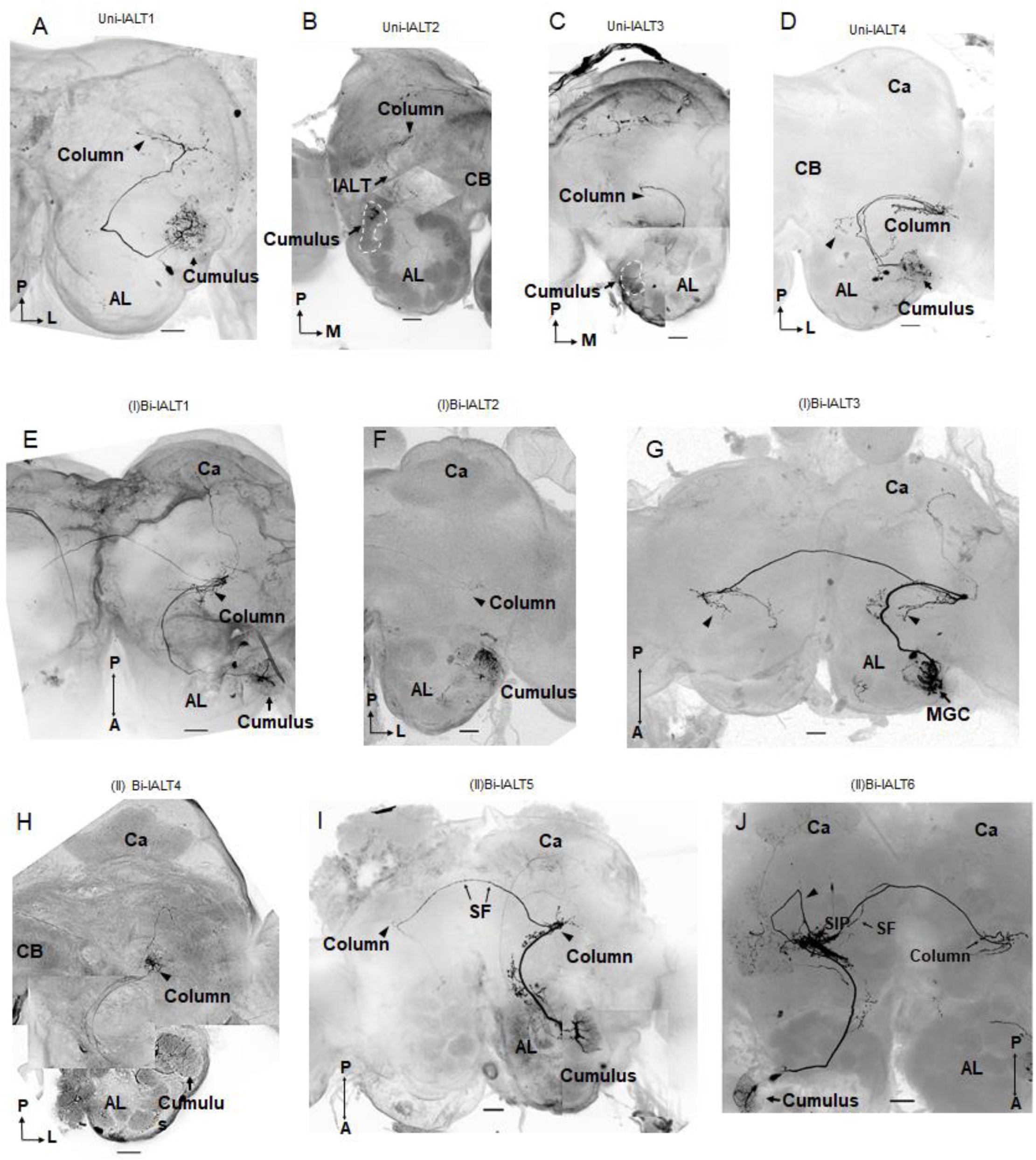
**Morphologies of all ten recorded lateral-tract neurons**. (**A-D**) Confocal images of the four unilateral lateral-tract neurons. (**E-G**) Confocal images of three Sub-type I bilateral lateral-tract neurons. (**H-J**) Confocal images of three Sub-type II bilateral lateral-tract neurons. AL, antennal lobe; Ca, Calyces of the mushroom body; CB, central body. A, anterior; L, lateral; M, medial; P, posterior; V, ventral. Scale bars, 50 μm.

**SM. Fig. 4.**
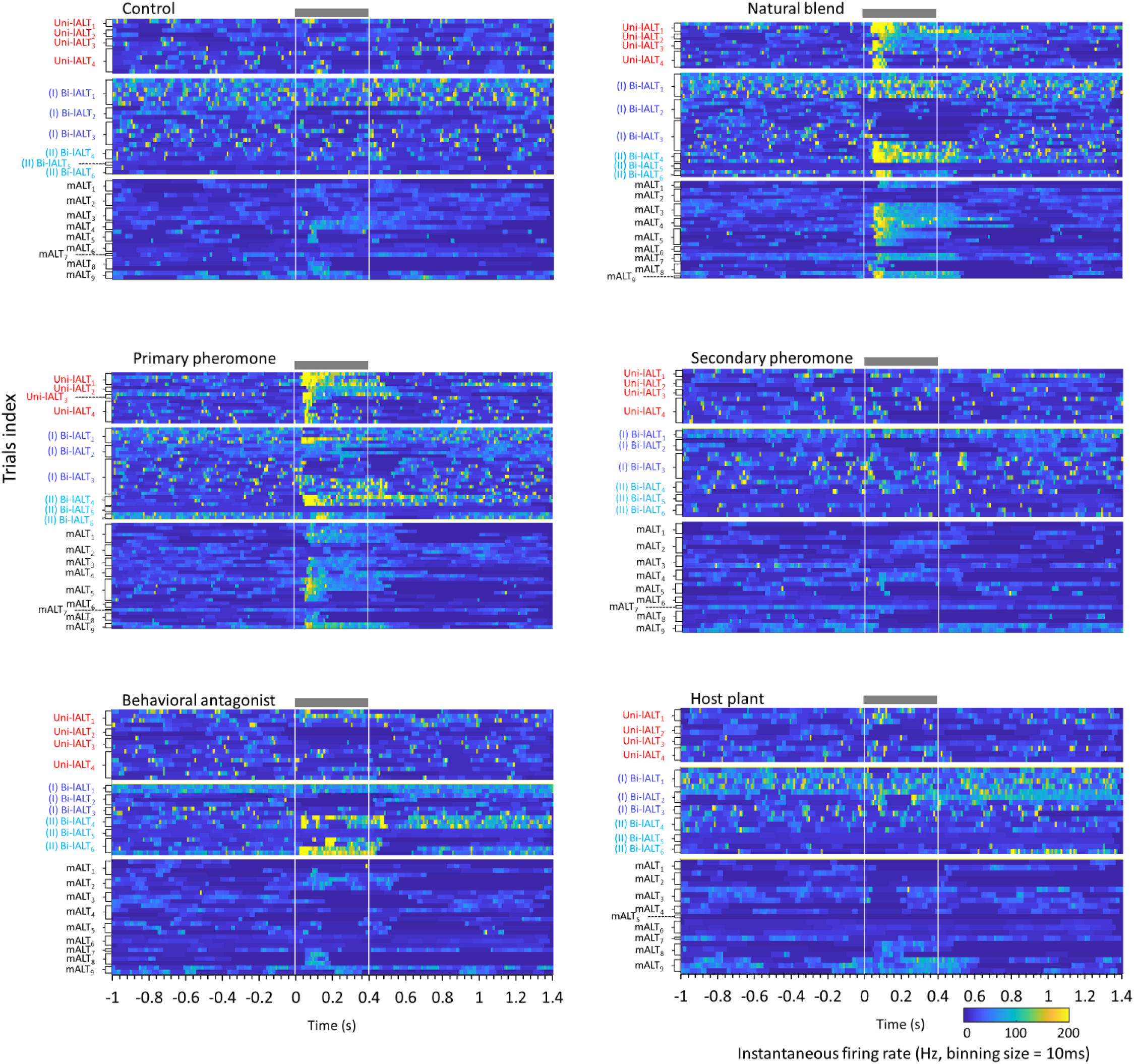
Neuron spiking activities of the individual antennal-lobe output neurons during application of pheromone and plant odor stimuli. Responding profiles can be seen for ten lateral-tract cumulus neurons and nine medial-tract cumulus neurons, illustrated by the instantaneous firing rate heat map. Each row plots an individual trial. The *grey* bar represents the onset and duration of the stimulus (400ms).

**SM. Fig. 5.**
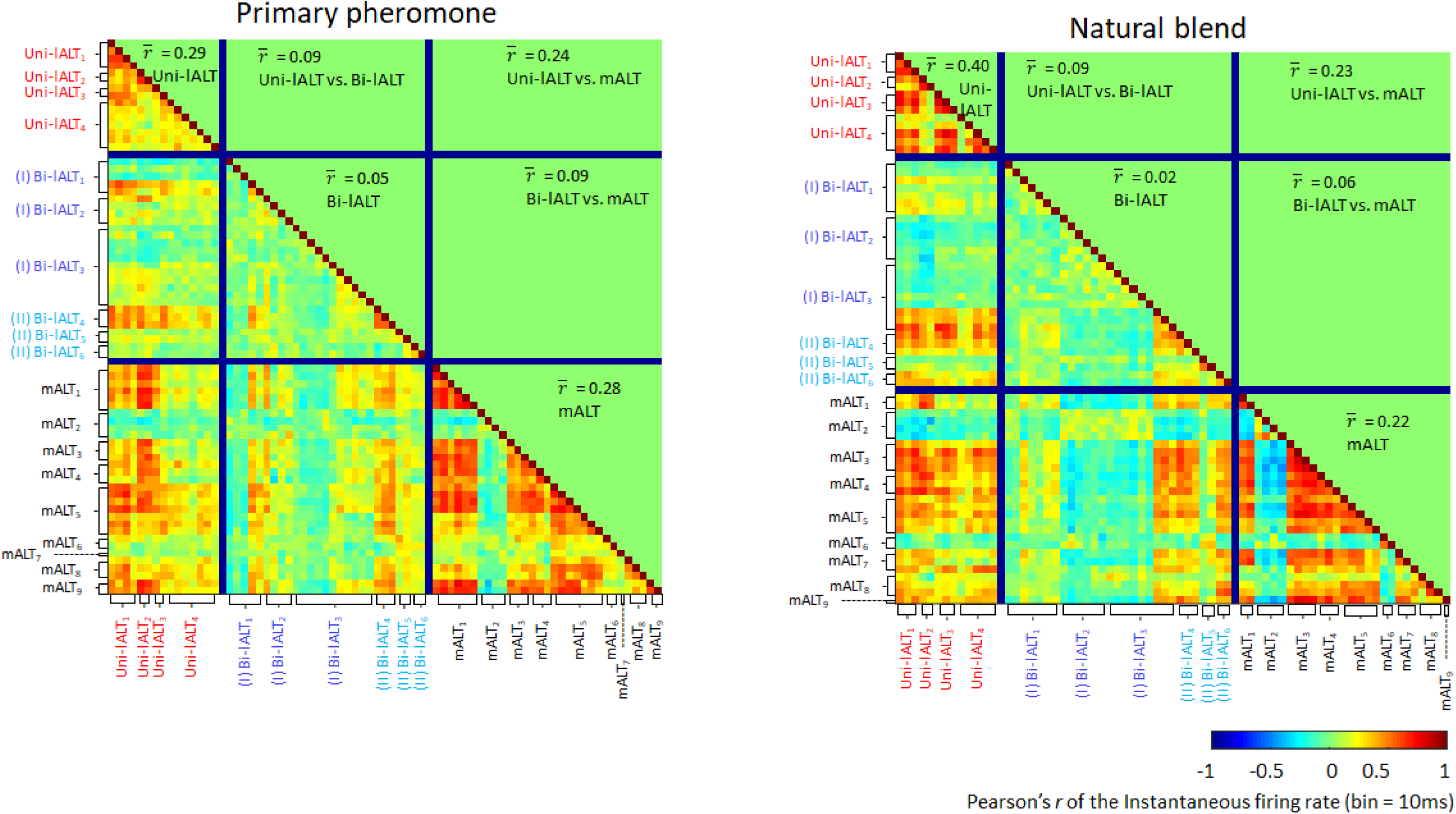
Pairwise correlation plots of the instantaneous firing rate (IFR, bin-size: 10 ms) to the primary pheromone (*left*) and natural blend (*right*). Each row plots the pearson’s *r* according to the IFRs of every two trials either within the same neuron type or across two different types. The mean correlation values 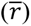 of neurons within each type or between two types are presented within the corresponding area located symmetric along the diagonal.

**SM. Fig. 6.**
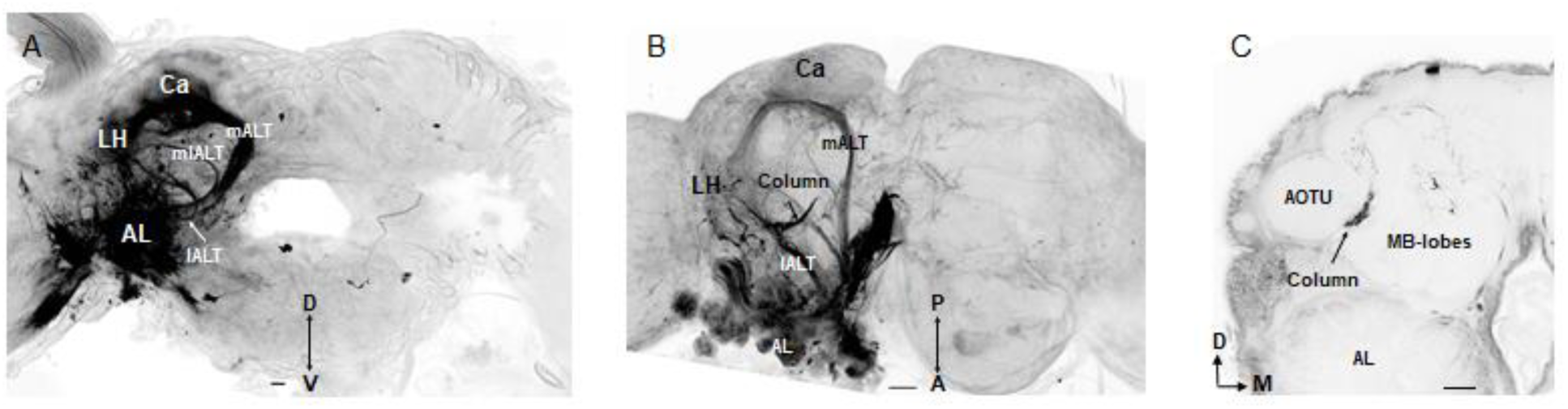
Projection profiles of antennal-lobe output neurons in other moth species. (**A**) Lateral-tract neurons in the Chinese oak silk moth, *Antheraea pernyi*, project to the lateral horn. (**B-C**) The column is the main target region of neurons confined to the lateral tract in the Silver Y, *Autographa gamma* (**B**) and Bogong moth, *Agrotis infusa* (**C**). Scale bars, 50 μm.

**SM. Fig. 7.**
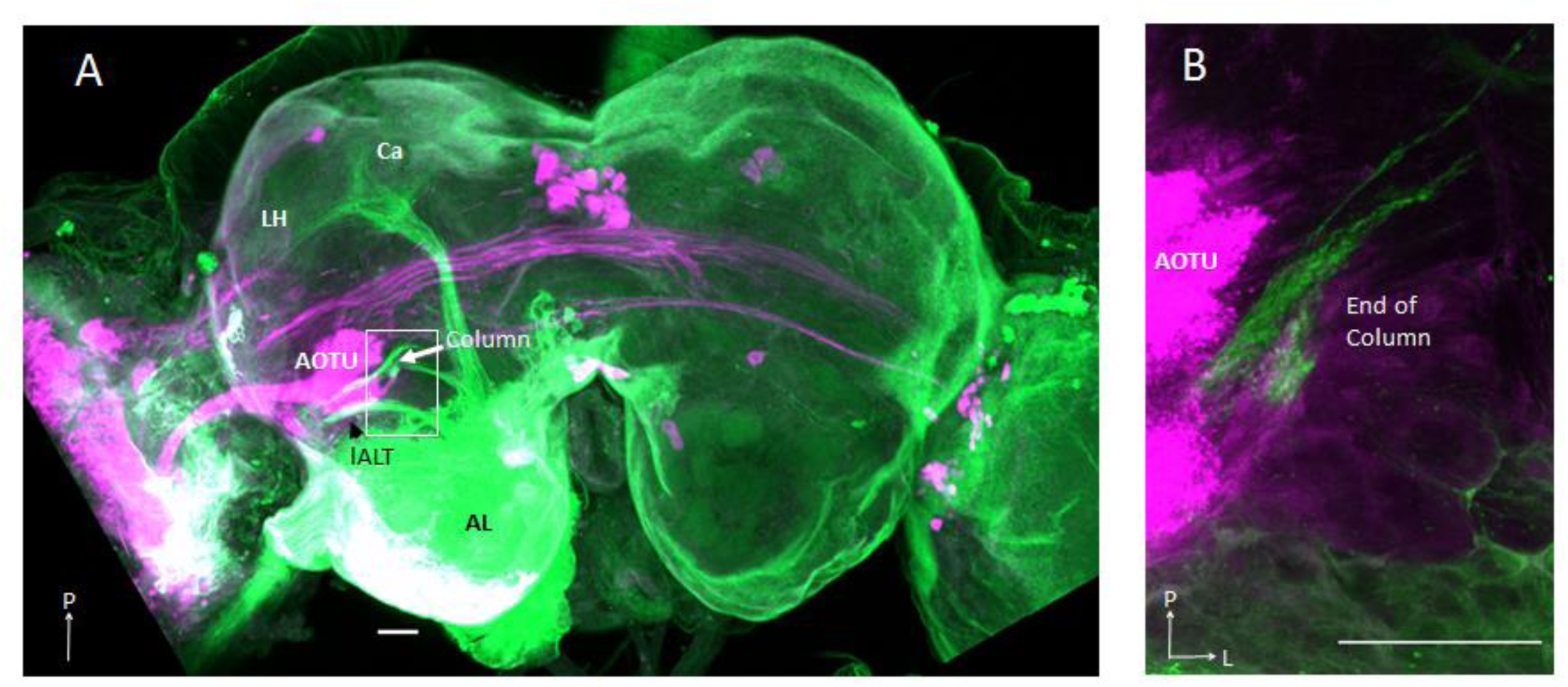
The target of antennal-lobe (AL) output neurons passing along the lateral tract is not overlapping with the prominent visual neuropil, the anterior optic tubercle (AOTU). (**A**) Confocal image in dorsal view of a double-labeled brain where different dyes were applied to the right optical lobe (*magenta*) and right antennal lobe (AL, *green*). (**B**) Magnified image of the area covered by the *white* square in (**A**), showing that the optic-lobe neurons projecting to the AOTU have no direct connection with the lateral-tract AL neurons. Scale bars, 50 μm.

**SM Fig.8.**
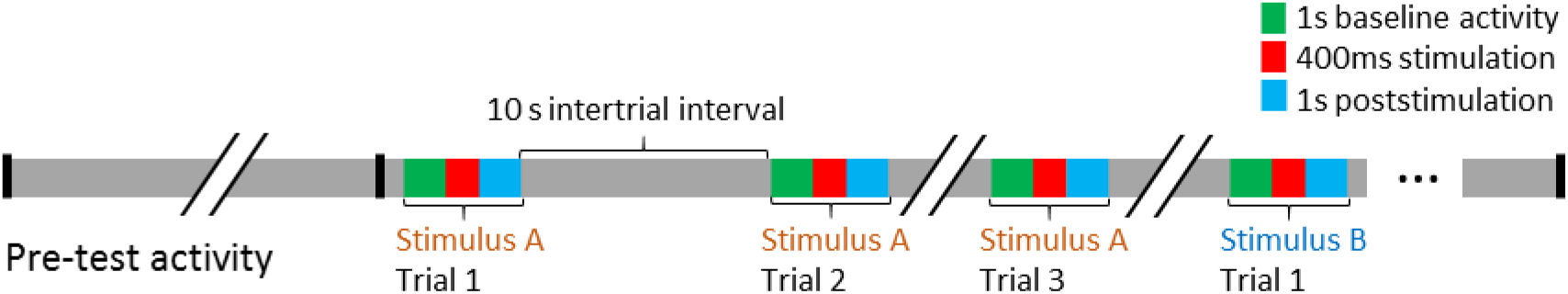
Experiment and stimulation protocol. After establishing stable contact with an individual neuron, spiking data was collected during a period of 25-40 seconds before stimulation. Each stimulus was tested at least two times, during a 400ms stimulation period, before iontophoretic staining. The analysis of spiking data includes 1 second baseline activity, 400ms stimulation, and 1 second poststimulation activity.

**SM Table 1.**
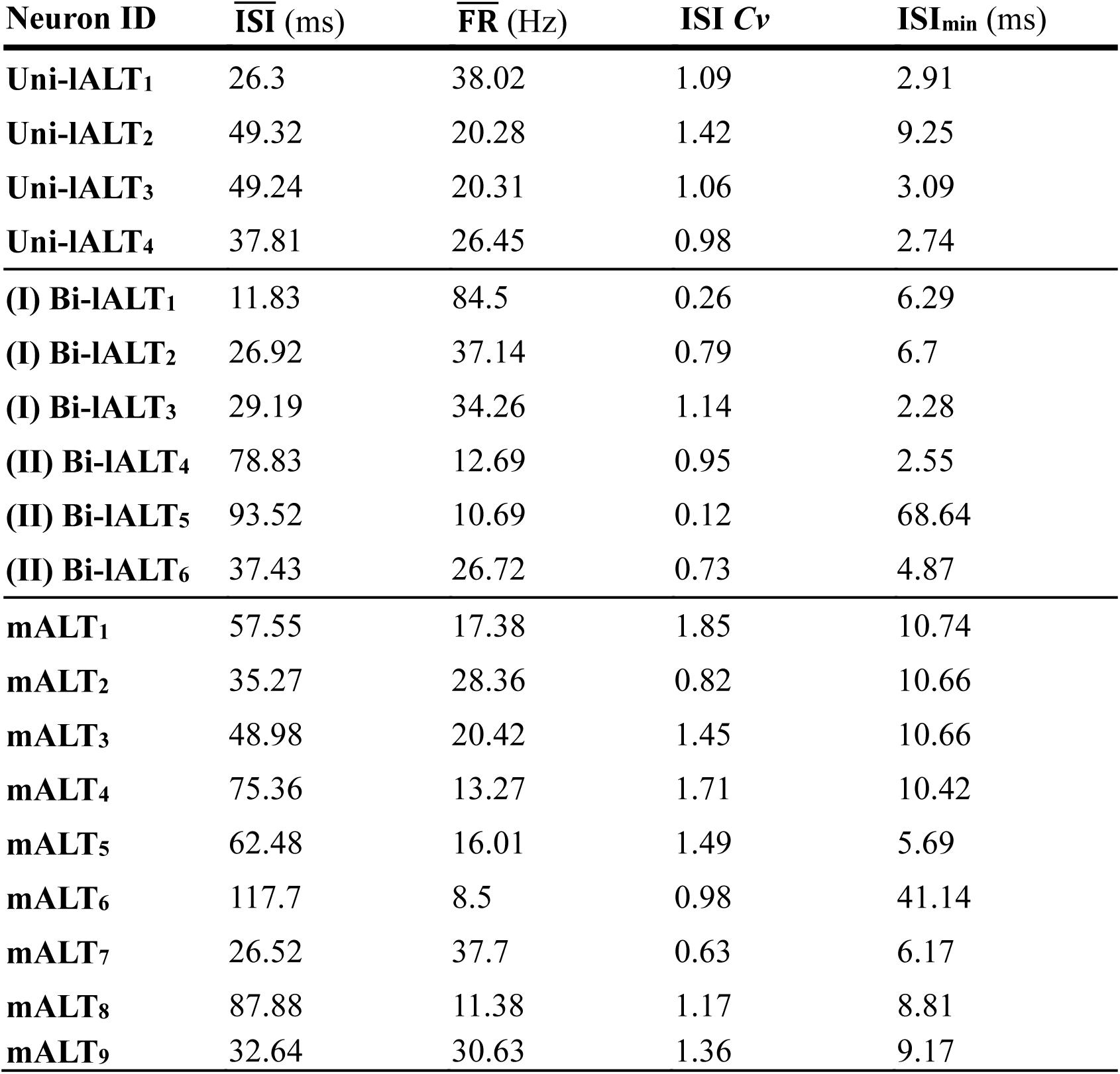
Overview of neuronal parameters during pre-test activity in recorded neurons confined to lateral tract and medial tract. 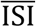 mean interspike interval, 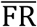: mean firing rate, ISI *Cv*: interspike interval coefficient of variation, ISI_min_: minimum interspike interval.

## References

Baker, T. C. & Hansson, B. S. 2016. Moth Sex Pheromone Olfaction. In: Allison, J. D. & CardŽ, R. T. (eds.) Pheromone Communication in Moths: Evolution, Behavior, and Application. Oakland: University of California Press.

Cho, S., Mitchell, A., Mitter, C., Regier, J., Matthews, M. & Robertson, R. 2008. Molecular phylogenetics of heliothine moths (Lepidoptera: Noctuidae: Heliothinae), with comments on the evolution of host range and pest status. Systematic Entomology, 33, 581–594.

Christensen, T. A. & Hildebrand, J. G. 1987. Male-specific, sex pheromone-selective projection neurons in the antennal lobes of the moth Manduca sexta. J Comp Physiol A, 160, 553–69.

Christensen, T. A., Mustaparta, H. & Hildebrand, J. G. 1991. Chemical communication in heliothine moths. Journal of Comparative Physiology A, 169, 259–274.

Christensen, T. A., Mustaparta, H. & Hildebrand, J. G. 1995. Chemical communication in heliothine moths. Journal of Comparative Physiology A, 177, 545–557.

Evers, J. F., Schmitt, S., Sibila, M. & Duch, C. 2005. Progress in functional neuroanatomy: precise automatic geometric reconstruction of neuronal morphology from confocal image stacks. Journal of Neurophysiology, 93, 2331–2342.

Fitt, G. P. 1989. The Ecology of Heliothis Species in Relation to Agroecosystems. Annual Review of Entomology, 34, 17–53.

Galizia, C. G. & Rossler, W. 2010. Parallel olfactory systems in insects: anatomy and function. Annu Rev Entomol, 55, 399–420.

Hamdani, E. H. & Døving, K. B. 2007. The functional organization of the fish olfactory system. Progress in Neurobiology, 82, 80–86.

Hansson, B. S., Christensen, T. A. & Hildebrand, J. G. 1991. Functionally distinct subdivisions of the macroglomerular complex in the antennal lobe of the male sphinx moth Manduca sexta. J Comp Neurol, 312, 264–78.

Homberg, U., Heinze, S., Pfeiffer, K., Kinoshita, M. & El Jundi, B. 2011. Central neural coding of sky polarization in insects. Philosophical transactions of the Royal Society of London. Series B, Biological sciences, 366, 680–687.

Homberg, U., Montague, R. A. & Hildebrand, J. G. 1988. Anatomy of antenno-cerebral pathways in the brain of the sphinx moth Manduca sexta. Cell and Tissue Research, 254, 255–281.

Ian, E., Kirkerud, N. H., Galizia, C. G. & Berg, B. G. 2017. Coincidence of pheromone and plant odor leads to sensory plasticity in the heliothine olfactory system. PloS one, 12, e0175513.

Ian, E., Zhao, X. C., Lande, A. & Berg, B. G. 2016. Individual Neurons Confined to Distinct Antennal-Lobe Tracts in the Heliothine Moth: Morphological Characteristics and Global Projection Patterns. Frontiers in Neuroanatomy, 10, 101.

Ito, K., Shinomiya, K., Ito, M., Armstrong, J. D., Boyan, G., Hartenstein, V., Harzsch, S., Heisenberg, M., Homberg, U., Jenett, A., Keshishian, H., Restifo, L. L., Rossler, W., Simpson, J. H., Strausfeld, N. J., Strauss, R. & Vosshall, L. B. 2014. A systematic nomenclature for the insect brain. Neuron, 81, 755–65.

Jacobs, G. A., Miller, J. P. & Aldworth, Z. 2008. Computational mechanisms of mechanosensory processing in the cricket. Journal of Experimental Biology, 211, 1819–1828.

Jarriault, D., Gadenne, C., Rospars, J.-P. & Anton, S. 2009. Quantitative analysis of sex-pheromone coding in the antennal lobe of the moth <EM=Agrotis ipsilon</EM=: a tool to study network plasticity. 212, 1191–1201.

Kanzaki, R., Arbas, E. A., Strausfeld, N. J. & Hildebrand, J. G. J. J. O. C. P. A. 1989. Physiology and morphology of projection neurons in the antennal lobe of the male mothManduca sexta. 165, 427–453.

Kanzaki, R., Soo, K., Seki, Y. & Wada, S. 2003. Projections to Higher Olfactory Centers from Subdivisions of the Antennal Lobe Macroglomerular Complex of the Male Silkmoth. Chemical Senses, 28, 113–130.

Kauer, J. S. 1991. Contributions of topography and parallel processing to odor coding in the vertebrate olfactory pathway. Trends in Neurosciences, 14, 79–85.

Kehat, M. & Dunkelblum, E. 1990. Behavioral responses of maleHeliothis armigera (Lepidoptera: Noctuidae) moths in a flight tunnel to combinations of components identified from female sex pheromone glands. Journal of Insect Behavior, 3, 75–83.

Kehat, M., Gothilf, S., Dunkelblum, E. & Greenberg, S. 1980. Field evaluation of female sex pheromone components of the cotton bollworm, Heliothis armigera. Entomologia Experimentalis et Applicata, 27, 188–193.

Kennedy, J. S. & Marsh, D. 1974. Pheromone-regulated anemotaxis in flying moths. Science, 184, 999–1001.

Kirschner, S., Kleineidam, C. J., Zube, C., Rybak, J., Grunewald, B. & Rossler, W. 2006. Dual olfactory pathway in the honeybee, Apis mellifera. J Comp Neurol, 499, 933–52.

Lee, S. G., Celestino, C. F., Stagg, J., Kleineidam, C. & Vickers, N. J. 2019. Moth pheromone-selective projection neurons with cell bodies in the antennal lobe lateral cluster exhibit diverse morphological and neurophysiological characteristics. J Comp Neurol.

Løfaldli, B., Kvello, P., Kirkerud, N. & Mustaparta, H. 2012. Activity in Neurons of a Putative Protocerebral Circuit Representing Information about a 10 Component Plant Odor Blend in Heliothis virescens. Frontiers in Systems Neuroscience, 6.

Mori, K. 2016. Chapter 1 - Axonal Projection of Olfactory Bulb Tufted and Mitral Cells to Olfactory Cortex A2 - Rockland, Kathleen S. Axons and Brain Architecture. San Diego: Academic Press.

Ott, S. R. 2008. Confocal microscopy in large insect brains: Zinc–formaldehyde fixation improves synapsin immunostaining and preservation of morphology in whole-mounts. Journal of Neuroscience Methods, 172, 220–230.

Patella, P. & Wilson, R. I. 2018. Functional Maps of Mechanosensory Features in the Drosophila Brain. Curr Biol, 28, 1189–1203.e5.

Pfeiffer, K. & Homberg, U. 2014. Organization and functional roles of the central complex in the insect brain. Annu Rev Entomol, 59, 165–84.

Piccardi, P., Capizzi, A., Cassani, G., Spinelli, P., Arsura, E. & Massardo, P. 1977. A sex pheromone component of the Old World bollworm Heliothis armigera. Journal of Insect Physiology, 23, 1443–1445.

Preibisch, S., Saalfeld, S. & Tomancak, P. 2009. Globally optimal stitching of tiled 3D microscopic image acquisitions. Bioinformatics, 25, 1463–1465.

Pryluk, R., Kfir, Y., Gelbard-Sagiv, H., Fried, I. & Paz, R. 2019. A Tradeoff in the Neural Code across Regions and Species. Cell, 176, 597–609.e18.

Raman, B., Ito, I. & Stopfer, M. 2008. Bilateral olfaction: two is better than one for navigation. Genome biology, 9, 212–212.

Rø, H., Müller, D. & Mustaparta, H. 2007. Anatomical organization of antennal lobe projection neurons in the moth Heliothis virescens. The Journal of Comparative Neurology, 500, 658–675.

Rodrigues, V. 1988. Spatial coding of olfactory information in the antennal lobe of Drosophila melanogaster. Brain Res, 453, 299–307.

Schmitt, S., Evers, J. F., Duch, C., Scholz, M. & Obermayer, K. 2004. New methods for the computer-assisted 3-D reconstruction of neurons from confocal image stacks. Neuroimage, 23, 1283–1298.

Seki, Y., Aonuma, H. & Kanzaki, R. 2005. Pheromone processing center in the protocerebrum of Bombyx mori revealed by nitric oxide-induced anti-cgmp immunocytochemistry. J Comp Neurol, 481, 340–51.

Skiri, H. T., Rø, H., Berg, B. G. & Mustaparta, H. 2005. Consistent organization of glomeruli in the antennal lobes of related species of heliothine moths. J Comp Neurol, 491, 367–80.

Strauch, M., Rein, J., Lutz, C. & Galizia, C. G. 2013. Signal extraction from movies of honeybee brain activity: the ImageBee plugin for Knime. Bmc Bioinformatics, 14, S4–S4.

Tanaka, N. K., Endo, K. & Ito, K. 2012. Organization of antennal lobe-associated neurons in adult Drosophila melanogaster brain. J Comp Neurol, 520, 4067–130.

Vickers, N. J. & Baker, T. C. 1994. Reiterative responses to single strands of odor promote sustained upwind flight and odor source location by moths. Proceedings of the National Academy of Sciences, 91, 5756–5760.

Vickers, N. J., Christensen, T. A. & Hildebrand, J. G. 1998. Combinatorial odor discrimination in the brain: attractive and antagonist odor blends are represented in distinct combinations of uniquely identifiable glomeruli. J Comp Neurol, 400, 35–56.

Willis, M. A., Avondet, J. L. & Zheng, E. 2011. The role of vision in odor-plume tracking by walking and flying insects. 214, 4121–4132.

Wu, D., Yan, Y. & Cui, J. 1997. Sex pheromone components of Helicoverpa armigera:chemical analysis and field tests. Insect Science, 4, 350–356.

Wu, H., Xu, M., Hou, C., Huang, L.-Q., Dong, J.-F. & Wang, C.-Z. 2015. Specific olfactory neurons and glomeruli are associated to differences in behavioral responses to pheromone components between two Helicoverpa species. Frontiers in Behavioral Neuroscience, 9, 206.

Wu, W., Anton, S., Lofstedt, C. & Hansson, B. S. 1996. Discrimination among pheromone component blends by interneurons in male antennal lobes of two populations of the turnip moth, Agrotis segetum. Proc Natl Acad Sci U S A, 93, 8022–7.

Zhao, X.-C. & Berg, B. G. 2010. Arrangement of Output Information from the 3 Macroglomerular Units in the Heliothine Moth Helicoverpa assulta: Morphological and Physiological Features of Male-Specific Projection Neurons. Chemical Senses, 35, 511–521.

Zhao, X.-C., Kvello, P., Løfaldli, B. B., Lillevoll, S. C., Mustaparta, H. & Berg, B. G. 2014. Representation of pheromones, interspecific signals, and plant odors in higher olfactory centers; mapping physiologically identified antennal-lobe projection neurons in the male heliothine moth. Frontiers in Systems Neuroscience, 8, 186.

Zhao, X. C., Chen, Q. Y., Guo, P., Xie, G. Y., Tang, Q. B., Guo, X. R. & Berg, B. G. 2016a. Glomerular identification in the antennal lobe of the male moth Helicoverpa armigera. J Comp Neurol, 524, 2993–3013.

Zhao, X. C., Ma, B. W., Berg, B. G., Xie, G. Y., Tang, Q. B. & Guo, X. R. 2016b. A global-wide search for sexual dimorphism of glomeruli in the antennal lobe of female and male Helicoverpa armigera. Scientific reports, 6.

